# piRAT: piRNA Annotation Tool for annotating, analyzing, and visualizing piRNAs

**DOI:** 10.1101/2025.07.29.667427

**Authors:** Dominik Robak, Guillem Ylla

## Abstract

**Motivation:** In recent years, Piwi-interacting RNAs (piRNAs) have been found to be involved in different biological roles beyond their initially identified role of protecting the germline genome. However, there is a lack of robust computational tools that facilitate their accurate annotation, analysis, and visualization, which would allow large-scale analysis of piRNAs across samples and species.

**Results:** We present piRAT, a **piR**NA **A**nnotation **T**ool for annotating both primary piRNAs from piRNA clusters and secondary piRNAs generated via the ping-pong cycle, using small RNA-seq data mapped to a genome. piRAT also performs descriptive analyses and generates comprehensive reports with visualizations to support annotations and provide further insights.

**Availability and Implementation:** piRAT is available on GitHub (https://github.com/ylla-lab/piRAT), implemented in Python 3, and released under MIT license. It is also installable through pip, conda, and available as Docker container (https://hub.docker.com/u/domrob). The Python implementation runs natively on Linux and in WSL in Windows, and the Docker image can run on Linux, Windows, and MacOS.

## 1. Introduction

Piwi-interacting RNAs (piRNAs) are a class of small non-coding RNAs, typically 26-33 nucleotides in length depending on the species, first identified in the early 2000s (Aravin *et al*. 2001). Similar to other small RNAs, such as micro RNAs (miRNAs) and silencing RNAs (siRNAs), piRNAs bind to Argonaute-family proteins, which mediate their interaction with target transcripts, regulating the target’s expression at a transcriptional or post-transcriptional level. In the case of piRNAs, as their name suggests, they bind to the PIWI proteins (Girard *et al*. 2006) a clade belonging to the Argonaute family (Carmell *et al*. 2002).

Initially, piRNAs were identified in germ cells of Drosophila, and subsequently in gonads of other animals, where they were shown to target transposable elements, a role for which they became known as guardians of the genome (Aravin, Hannon and Brennecke 2007). However, more recent studies have revealed that piRNAs are also abundantly expressed in both gonadal and non-gonadal somatic tissues in many animal species (Lewis *et al*. 2018; Llonga *et al*. 2018; Yamashita *et al*. 2024). These discoveries have broadened the focus towards the roles of piRNAs beyond the germline. Indeed, piRNAs have been found to be involved in a variety of biological processes, including sex determination in silkworms (Kiuchi *et al*. 2014), multiple human cancers (Liu *et al*. 2019; Weng, Li and Goel 2019; Chen *et al*. 2021), and early embryo development (Ramat and Simonelig 2021).

Despite their important roles, there is a limited amount of available tools to robustly annotate piRNAs, which is a crucial step for studying them. This lack of tools is especially detrimental for research on non-model organisms for which no prior small RNA annotations, including piRNA annotations and piRNA databases, exist (Montañés *et al*. 2021; Mito *et al*. 2022; Nakamura, Ylla and Extavour 2022). In this article, we present **piRAT**, a **piR**NA **A**nnotation **T**ool, that leverages small RNA-seq data aligned to a reference genome to annotate piRNA regions from the primary and secondary pathway based on information regarding the piRNA biogenesis, suitable for both model and non-model organisms.

### 1.1 piRNA biogenesis signatures to annotate piRNAs

The piRNA biogenesis is most well understood in the model organism *D. melanogaster*, and although some steps are still under discussion, most animals share similar mechanisms of piRNA biogenesis (Gainetdinov *et al*. 2018). It is well established that piRNAs can be generated through two pathways, the primary pathway and the secondary pathway, also known as the ping-pong cycle. Both pathways display specific signatures in the small RNA-seq data that piRAT leverages to annotate bona fide piRNAs loci.

In brief, the primary pathway starts with the transcription of long primary transcripts (up to hundreds of kilobases long) that are subsequently cleaved by Zucchini protein, releasing small RNA fragments (pre-piRNAs). These RNAs are loaded onto a PIWI protein, followed by 3’ trimming and 2-O-methylation, resulting in the 26-33 nts long mature primary piRNAs (Gainetdinov *et al*. 2018). Based on this knowledge, we can identify regions producing primary transcripts, also known as piRNA clusters, as genomic regions with a high density of small RNA-seq reads of the length corresponding to the piRNAs on the given species.

The secondary pathway, known as the ping-pong cycle, was originally discovered in *D. melanogaster* germline (Brennecke *et al*. 2007; Gunawardane *et al*. 2007) and subsequently detected in other animals’ germline, embryogenesis, and more recently in non-gonadal tissues of some animal lineages (Houwing, Berezikov and Ketting 2008; Kawaoka *et al*. 2008, Kawaoka *et al*. 2011; Yamashita *et al*. 2024). In this cycle, piRNAs bind to PIWI (or its paralogue in flies Aubergine) and identify their target RNAs based on sequence complementarity. Vasa and Krimper cleave the target at the position corresponding to the complementary 10^th^ nucleotide of the primary piRNA. Then, AGO3 binds to the 5’of the cleaved RNA and Nibbler trims the 3’ end, leaving ∼28nts sequence bound to AGO3, which is then methylated by Hen1, producing a mature secondary piRNA. This mature secondary piRNA, is complementary to the piRNA cluster transcripts, which it slices, producing more primary piRNAs as the one that initiated the cycle (more details in our recent publication (Yamashita *et al*. 2024)).

The presence of the ping-pong cycle can be robustly determined through small RNA-seq pathway by the so-called ping-pong signature (Czech and Hannon 2016). This ping-pong signature consists of the two small RNA-seq reads of ∼28 nts with a 5’ to 5’ overlap of 10 nts in opposite strands of the genome. Additionally, because primary piRNAs are enriched with a U at the 5’ end, secondary piRNAs are enriched by an A at position 10.

### 1.2 Tools for piRNA annotation and analysis

Despite the relevance of piRNAs, few methods exist for robust identification, annotation, classification, and visualization of piRNAs that can work across different animals.

Additionally, the few existing tools present significant limitations (**Table 1**).

**Table 1:** Feature comparison of the principal piRNA annotation tools and the herein presented piRAT. Comparison includes whether they can annotate piRNA clusters and ping-pong loci, the required type of inputs and their format, the generation of graphical outputs, and the use of multiple CPU threads.

|  | Annotation of piRNA clusters | Annotation of ping-pong loci | Input file type | No special pre-processing | Graphical output | Max. number of input files per run | Multi-thread computing |
| --- | --- | --- | --- | --- | --- | --- | --- |
| piRAT | ✓ | ✓ | .bam | ✓ | ✓ | Limited by hardware | ✓ |
| proTRAC | ✓ | ✗ | .sam | ✗ | ✓ | 1 | ✗ |
| piClust | ✓ | ✗ | .sam | ✗ | ✗ | 1 | ✗ |
| PILFER | ✓ | ✗ | .bed | ✗ | ✗ | 1 | ✗ |
| PingPongPro | ✗ | ✓ | .sam/.bam | ✗ | ✓ | 1 | ✗ |

Among the available tools for piRNA cluster annotation, proTRAC (Rosenkranz and Zischler 2012), is one of the most widely used. However, while proTRAC specializes in piRNA cluster annotation, it does not annotate ping-pong piRNAs. Furthermore, due to its probabilistic method based on the assumption of uniform distribution, proTRAC was found to fail to accurately annotate piRNA clusters in some species (Jung, Park and Kim 2014). In response to this limitation, another tool called piClust (Jung, Park and Kim 2014) was developed based on the known Density-Based Algorithm for Discovering Clusters in Large Spatial Databases with Noise (DBSCAN) (Ester *et al*. 1996).

While piClust was shown to outperform proTRAC in piRNA cluster discovery (Jung, Park and Kim 2014), it has several limitations that make it unsuitable for many purposes. A major limitation is that piClust ran through a web server that accepted a single SAM file with a maximum size of 200Mb per submission. Additionally, at the time of writing, the piClust server is down. A further drawback of piClust and proTRAC, is the lack of ping-pong annotations, forcing the user to look for alternative methods to identify ping-pong sites.

One of the most recently published tools for piRNA cluster annotation is PILFER (Ray and Pandey 2018). When the authors benchmarked it against proTRAC, PILFER identified fewer clusters but longer (Ray and Pandey 2018). However, PILFER was not designed to work solely with small RNA-seq reads mapped to a genome. In their publication (Ray and Pandey 2018), the authors used human small RNA-seq reads that were either classified as canonical piRNAs or potential piRNAs. To identify canonical piRNA reads, they mapped the reads to the human genome and retained those overlapping with piRNAs annotated in piRNABank. Potential piRNAs were defined as 26–33 nucleotide reads that mapped to the genome but were not found in piRNABank and did not overlap with other known non-coding RNAs. The final input for PILFER was a BED file containing the genomic coordinates of both canonical and putative piRNAs. This pre-processing requires extra work and relies heavily on prior piRNA and non-coding RNA annotations, limiting its utility in non-model species.

Our tool, **piRAT**, addresses the limitations of existing piRNA annotation tools and offers several novel features, including comprehensive graphical outputs that visually support the annotations. Consequently, piRAT stands out for **a)** identifying piRNAs from both biogenesis pathways in a single run, **b)** accepting multiple standard binary alignment map (BAM) files of any size (with limits determined by hardware capabilities), **c)** without requiring special pre-processing steps or need for external databases or annotations), **d)** generating graphical outputs that support the annotations and provide overall descriptive statistics.

Thus, piRAT provides a comprehensive set of features, utilizes standard input and output file formats, and doesn’t require any type of genome annotations. Moreover, as demonstrated subsequently, it also yields higher-quality piRNA annotations compared to existing tools.

## 2 Methods and implementation

As input, piRAT requires a directory containing BAM files of small RNA-seq aligned to the same reference genome. The output consists of general feature format (GFF) files with the genomic coordinates of the annotated piRNA clusters and ping-pong signature sites found within the samples. Additionally, piRAT returns comprehensive graphical reports displaying the supporting data for the annotations and other summary statistics.

While **piRAT** executes with a single command, internally it comprises four distinct modules: the first module determines the piRNA length within the provided dataset; the second module annotates the primary piRNA clusters; the third module identifies the ping-pong loci; and the fourth module generates graphical reports of both the input data and final annotations **(Figure 1**). Each of these modules is described in detail below.

**Figure 1:**
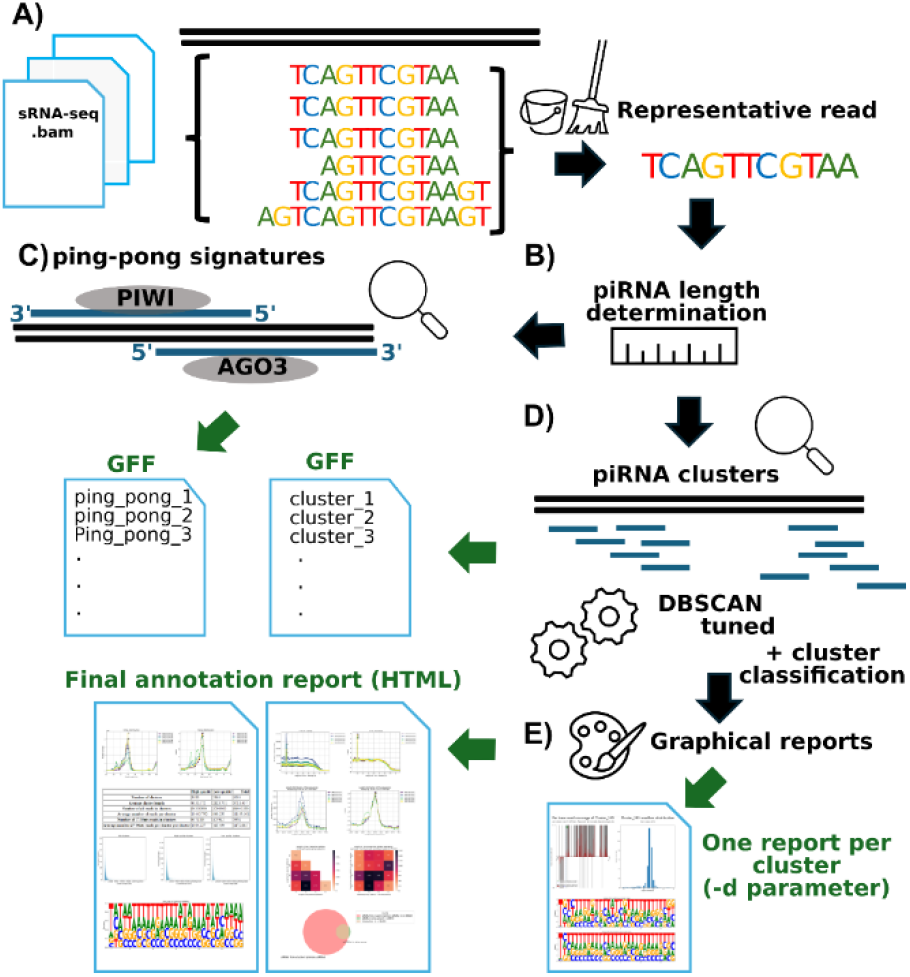
Schematic workflow of piRAT. **A)** Starting with small RNA-seq datasets mapped to a reference genome, piRAT first cleans the data by selecting a single representative read per locus. **B)** Then determines the piRNA length in the dataset (if not provided by the user), and proceeds with the **C)** annotation of ping-pong loci using the characteristic ping-pong signature, and **D)** piRNA clusters using a tuned version of DBSCAN followed by a cluster classification step. Finally, **E)** piRAT generates extensive graphical outputs in the final report, and optionally, one report per cluster with plots showing its supporting data. The outputs (green arrows) are GFF files with annotations and HTML files with reports.

### 2.1 piRNA length detection and read variant purging

The piRNAs often range from ∼26-32 nts, but the specific lengths depend on the species. If the length of piRNAs in a particular species is known, the user can manually input this information into piRAT as a parameter. Alternatively, piRAT predicts the range of piRNA sizes for a given species based on the length of reads displaying 10 nt 5’ to 5’ overlap, or in the small RNA-seq read length distribution when running in the cluster-only mode (**Supplementary Methods 1**). Only reads that fall within the specified piRNA lengths, which can be either provided by the user or determined by piRAT, will be utilized in subsequent piRNA annotation steps. Reads within this length range are herein referred to as “piRNA-length reads”.

In small RNA-seq data, reads mapped in the same genomic locus might be slightly different even if they come from the same genomic element (miRNA, piRNA, etc.) (Raabe *et al*. 2014). This can be due to technical reasons, like errors during sequencing, or biological phenomena such as 3’-tailing, presence of immature RNA molecules, degradation, polymorphisms, and other post-transcriptional modifications. Often, we observe that for a given miRNA or piRNA, there is one read with a large number of copies and several other reads with slight variation in the extremes (mostly in 3’) with much fewer copies.

We argue that treating all unique small RNA-seq reads as distinct RNA molecules artificially inflates the number of annotated piRNAs. To mitigate this issue, piRAT selects a single “representative read” for each locus where multiple piRNA-length reads are mapped. The representative read is defined as the highest copy number among those mapped reads that exhibit up to three nucleotide differences at the start and end positions. The variation threshold is set to three nucleotides by default, based on our analysis, which showed that beyond three, the number of ping-pong signatures plateaus (**Supplementary Figure 1**). Users can manually adjust this threshold using the parameter (`--variation-threshold, -vt`), with a minimum value of zero, which would force piRAT to include all mapped reads with unique start and end coordinates.

### 2.2 Primary piRNA cluster annotation

In the primary piRNA biogenesis pathway, long precursor RNAs are cleaved into shorter fragments, bound to PIWI proteins, and trimmed, producing mature piRNAs. When small RNA-seq reads from these piRNAs are mapped to the genome, the reads originating from the same precursor RNA align to the genomic region of the precursor, typically referred to as a primary piRNA cluster. Annotating these piRNA clusters is the standard method for identifying piRNAs produced through the primary pathway (Rosenkranz and Zischler 2012; Jung, Park and Kim 2014; Ray and Pandey 2018).

To identify piRNA clusters, piRAT, similarly to piClust, is based on DBSCAN algorithm (Ester *et al*. 1996), although with some modifications explained below. DBSCAN requires two parameters k and eps, with k parameter being the k-th nearest neighbor used to compute the distance, and eps defining the maximum distance between two points (in this case, two small RNA-seq reads) for one to be considered within the neighborhood of the other. If the k-th neighbor is within eps distance, it means that the two points are connected by density-reachable, and hence belong to the same cluster (more details in **Supplementary Methods 2**). Both k and eps, can be provided by the user. If not specified, piRAT will estimate the optimal values for these parameters as described in **Supplementary Methods**.

Since piRAT’s cluster detection is strand-specific, an additional step is performed to merge clusters into bi-directional or dual-stranded. In this step, piRAT measures the distance from each cluster to its neighbors in the complementary strand, and if the distance between the 3’ ends is within the Eps value used during clustering, they are merged into a bi-directional cluster. If the length of overlap is greater than 50% of the shorter cluster, they are merged into a dual-strand cluster (**Supplementary Methods 2**).

Finally, piRAT classifies clusters as high- or low-quality based on specific biogenesis criteria. Using all small RNA-seq reads mapped within an annotated cluster, piRAT assigns a high-quality classification if two conditions are met: **(1)** more than 50% of the reads start with uridine (T, in the RNA-seq data), and **(2)** over 50% of the reads mapped within the cluster are of the defined piRNA length. Both thresholds can be adjusted by the user.

The cluster annotations are provided to the user in three GFF files: one containing all clusters, one for high-quality clusters, and one for low-quality clusters. Although both high- and low-quality clusters are returned, we recommend using only the high-quality clusters, as these are used in the subsequent benchmarking analyses. Additional output files are also generated, including information such as the genomic coordinates of all putative piRNAs within the clusters.

### 2.3 Ping-pong piRNA annotation

To identify piRNAs generated through the ping-pong cycle, piRAT utilizes the well-established “ping-pong signature” (Czech and Hannon 2016). This signature is characterized by pairs of piRNA-length small RNA-seq reads mapped to opposite strands of the genome, with a 5’ to 5’ overlap of 10 nucleotides. piRAT detects this ping-pong signature using the “representative reads” of piRNA length. The resulting ping-pong site annotations are provided in a GFF file containing the coordinates of these two pairs of potential piRNAs.

This GFF can be used downstream to quantify the expression of each specific ping-pong pair across samples and can also be visualized in genome browsers like IGV.

### 2.4 Graphical reports

The graphical outputs generated by piRAT are designed to provide support for the annotations and give an overview of the characteristics of the annotated piRNAs in a given dataset.

When piRAT runs with the active graphical output for clusters parameter (-d), it generates several plots for each cluster, including a bar plot showing the per-nucleotide read coverage along the cluster, the read length distribution, and two sequence logos representing all reads within the cluster and the other displaying only the representative reads. Although only piRNA-length representative reads are used for annotating clusters, all reads mapped to the cluster are included in these plots to provide comprehensive evidence of cluster quality. Since generating these graphical outputs increases runtime and hardware requirements when hundreds of clusters are annotated, this option is disabled by default.

In addition to the individual plots for each cluster, at the end of the run, piRAT generates the “piRAT Final Report”, which includes the following plots and tables (**Figure 2**):

**Figure 2:**
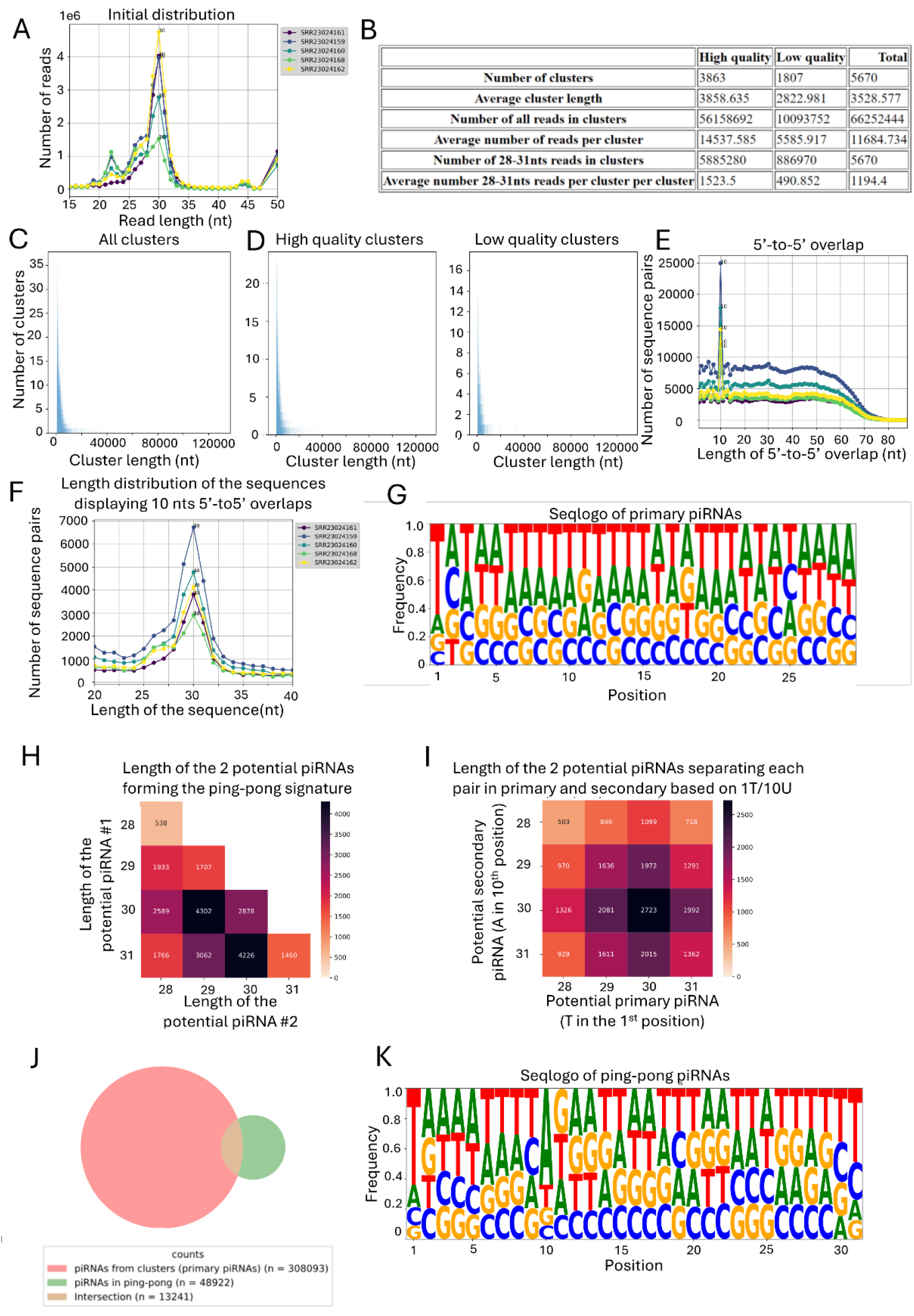
Example plots of the graphical output of piRAT running on 5 small RNA-seq datasets. **A)** Read length distribution in each dataset. **B)** Summary of the annotated clusters. **C)** Distribution of all the annotated clusters, as well as only for the **D)** high-quality and low-quality ones. **E)** 5’ to 5’ overlaps in each of the datasets. **F)** Read length distribution of those read pairs that have 10nts overlap 5’ to 5’. **G)** Seqlogo of the reads belonging to clusters. **H)** Heatmap of the piRNA length of the read pairs displaying 10nts 5’to 5’ overlap, as well as **I)** separating the pairs based on the ones starting with T and those with A at the 10^th^ position. **I)** Venn diagram displaying the overlap of piRNAs found in clusters as well as in ping-pong sites. **K)** SeqLogo of all the ping-pong piRNAs.

#### Initial Read Length Distribution Plot

Shows the distribution of small RNA-seq reads (20-40nts) across samples, with the x-axis representing read length and the y-axis the read count (**Figure 2A**). A z-score normalized version is also provided to compare samples with differing library sizes. The presence and height of a peak around 26-33 nt, provide clues about the relative abundance of piRNAs versus other small RNAs in the samples and length of the piRNAs in the given species.

#### Table of piRNA Cluster Statistics

This table summarizes key statistics for high-quality, low-quality, and total annotated piRNA clusters (**Figure 2B**).

#### Cluster Length Distribution

These three plots illustrate the length distributions of high-quality, low-quality, and all clusters(**Figure 2C-D**).

#### Primary piRNA Sequence Logo

This plot displays the sequence logo of reads inside the annotated clusters that are within the piRNA length range. Given the known enrichment of uracil at the first position of primary piRNAs, bona fide piRNA clusters should exhibit a clear bias toward T in this position **(Figure 2G)**.

#### 5’ to 5’ Overlaps plot

It displays the number of read pairs for each 5’ to 5’ overlaps value. A clear peak at 10 nts overlap is a strong indication of the presence of ping-pong piRNAs in the given sample **(Figure 2E)**. A z-score normalized version of the plot is also provided for comparing the intensity of the ping-pong signature across samples.

#### Length distribution of ping-pong signature sequence pairs

This plot displays the length distributions of the read pairs exhibiting a 10nt 5’ to 5’ overlap on opposite strands. It is enriched for true ping-pong piRNAs, providing clearer insights into the actual lengths of piRNAs involved in the ping-pong cycle for each sample. A z-score normalized version is also included. Representative reads within the piRNA-length range and showing this overlap are considered piRNAs participating in the ping-pong cycle **(Figure 2F)**.

#### Heatmap of Potential piRNA Pairs Lengths

This heatmap shows the frequencies of the lengths of the two piRNA displaying ping-pong signatures **(Figure 2H)**. This allows the user to determine whether there is homotypic ping-pong (the two piRNAs are of the same length) or heterotypic ping-pong (between piRNAs of different lengths).

#### Heatmap of Potential piRNA Pair Lengths Split by 1T and 10A

This heatmap differs from the previous on having on one axis the putative primary piRNAs displaying 1T and the putative secondary piRNA with 10A on the other axis **(Figure 2I)**. In the case of heterotypic ping-pong, this plot allows extrapolating what piRNA lengths are generated from the primary and secondary pathways.

#### Venn diagram

Displays the number of piRNAs in clusters and ping-pong sites, and the intersection shows the number of piRNAs found in a cluster that also participate in the ping-pong in other loci **(Figure 2J)**.

#### Sequence logo of ping-pong piRNAs

It contains the sequences of all representative reads of the detected ping-pong loci **(Figure 2K)**. Primary piRNAs typically enrich for U in the first position, while secondary piRNAs are enriched for A in the tenth position. If ping-pong piRNAs have been correctly identified, this sequence logo should show a higher frequency of T in the first position and a higher frequency of A at the tenth position.

## 3 Results

We benchmarked piRAT using publicly available small RNA-seq datasets from testes, ovaries, and non-gonadal tissue from humans (*Homo sapiens*), mice (*Mus musculus*), and flies (*Drosophila melanogaster*) (**Supplementary Table S1**). Male and female gonadal tissues were expected to exhibit piRNA expression, while somatic tissues served as negative controls, as piRNA expression in mammals is primarily restricted to germ cells. In flies, non-gonadal piRNA expression is possible albeit much reduced compared to gonads (Yamashita *et al*. 2024).

### 3.1 Ping-Pong signature annotation benchmark

We compared piRAT’s ping-pong signature annotations to those of the PingPongPro (Uhrig and Klein 2019) using the same small RNA-seq datasets from human, mouse, and fly testes and ovaries (**Table 2**). While piRAT provides coordinates for pairs of piRNAs displaying the signature along with comprehensive graphical plots, PingPongPro only returns the regions where the 10nt overlap occurs. In this comparison, piRAT identified approximately 42-67% of the signatures reported by PingPongPro.

**Table 2:** Number of identified ping-pong signatures by piRAT and PingPongPro in small RNA-seq data from testes and ovaries of humans, mice, and flies, and percentage of overlapping signatures between tools.

|  |  | piRAT | PingPongPro | % piRAT in pingPongPro | % pingPongPro in piRAT |
| --- | --- | --- | --- | --- | --- |
| **Human** |  |  |  |  |  |
|  | **Testes** | 60,457 | 174,441 | 100% | 58.32% |
|  | **Ovaries** | 9,314 | 13,641 | 100% | 56.65% |
| **Mouse** |  |  |  |  |  |
|  | **Testes** | 22,385 | 22,603 | 100% | 42.93% |
|  | **Ovaries** | 14,036 | 43,781 | 100% | 48.74% |
| **Fly** |  |  |  |  |  |
|  | **Testes** | 52,881 | 72,219 | 92% | 67.36% |
|  | **Ovaries** | 139,824 | 227,094 | 100% | 61.57% |

This discrepancy arises from piRAT’s stringent criteria to reduce false positives. While PingPongPro utilizes all unique reads, piRAT considers only reads of the specified piRNA length and utilizes a single representative read per locus.

### 3.2 piRNA cluster identification with real data

For piRNA clustering annotation, we compared piRAT to the other popular tools, proTRAC, and PILFER, using the same datasets of human, mouse, and fly from gonadal and non-gonadal tissue without any specific pre-processing **(Table 3**). Note that we were unable to include piClust in our benchmark due to the input file limit and the availability of the web server.

**Table 3:** Number of predicted clusters in each small RNA-seq datasets from testes, ovaries, and somatic tissues of humans, mice, and flies.

|  | Tissue | piRAT High-quality | piRAT Low-quality | proTRAC | PILFER |
| --- | --- | --- | --- | --- | --- |
| **Human** |  |  |  |  |  |
|  | **Testes** | 22,552 | 1,332 | 174 | 174 |
|  | **Ovaries** | 3,978 | 1,157 | 159 | 244 |
|  | **Somatic** | 0 | 28 | 0 | 5 |
| **Mouse** |  |  |  |  |  |
|  | **Testes** | 4,237 | 3,400 | 207 | 377 |
|  | **Ovaries** | 6,591 | 4,182 | 8 | 56 |
|  | **Somatic** | 0 | 59 | 0 | 95 |
| **Fly** |  |  |  |  |  |
|  | **Testes** | 2,630 | 3,038 | 6 | 135 |
|  | **Ovaries** | 4,430 | 395 | 5 | 1426 |
|  | **Somatic** | 9 | 1,807 | 0 | 47 |

piRAT consistently annotates a higher number of high-quality clusters than proTRAC and PILFER in gonadal tissues (**Table 3**). Considering that in our negative controls (the mammalian somatic tissues), no high-quality clusters are annotated by piRAT, the elevated number of high-quality clusters annotated by piRAT in gonadal tissues is unlikely to be due to false positives. While proTRAC also didn’t annotate any clusters in the mammalian non-gonadal tissues, PILFER found 5 in humans and 95 in mice (**Table 3)**. The clusters detected in *D. melanogaster*, are not necessarily false positives, given that non-gonadal piRNAs can occur in this species (Yamashita *et al*. 2024) and that the further explorations of the 9 clusters piRAT detected display clear signs of bona fide piRNA clusters **(Supplementary Figure S2**).

As an indicator of the cluster’s quality, we calculated the proportion of reads within the clusters annotated from each tool that are putative piRNAs (those of 26-32nts that start with T), assuming that real piRNA clusters should contain most of the reads having these characteristics (**Supplementary Table S2**). Here, piRAT high-quality clusters contained between 43.79% of 65.91% of reads with piRNA characteristics, similarly to proTRAC (51.50%-69.07%), and higher than PILFER (0.97%-47.78%).

As an indicator of the completeness of the piRNA clusters of each tool, we calculated the proportion of putative piRNA reads (those between 26-32 nts that start with T) within the RNA-seq datasets that map inside the clusters. Highly complete piRNA cluster annotations should contain a large fraction of these reads, assuming that most reads of these characteristics should belong to primary piRNAs and therefore should be found inside clusters (**Supplementary Table S3**). In this metric, piRAT consistently scores higher, with its high-quality clusters containing between 33.90% and 93.04% of the reads of piRNA characteristics, compared to proTRAC (0.24-53.12%) and PILFER (0.51%-56.53%).

Overall, piRAT annotated more clusters across species in gonadal tissues without showing signs of false positives, as it did not detect clusters in mammalian somatic tissues. piRAT high-quality clusters contain a high percentage of the small RNA-seq reads of the piRNA characteristics, and additionally, these clusters explain a higher proportion of small RNA-seq reads of piRNA characteristics.

### 3.3 Benchmarking against references

Due to the lack of a definitive ground truth for piRNA cluster annotations, we compared the performance of piRNA annotation tools against two different sources. First, we compare the cluster to those from mouse and human, annotated in Girard et al. (2006), which is the same dataset used by the authors of piClust for their benchmarks (Jung, Park and Kim 2014). The second benchmark was done using piRNAdb (https://www.pirnadb.org) (Piuco and Galante 2021), which contains clusters identified using the method proposed by (Lau *et al*. 2006) and using described piRNAs from multiple works.

We computed the percentage of bases in the reference clusters (from Girard et al. (2006) and piRNAdb) that overlap with the annotated clusters from piRAT, proTRAC, and PILFER (**Table 4**). In these comparisons, piRAT high-quality clusters in most cases consistently covered a higher percentage of the reference clusters than proTRAC and PILFER, indicating higher sensitivity.

**Table 4:** Percentage of nucleotides of piRNA clusters from piRNAdb and Girard. et al. 2006, that are covered by clusters annotated by piRAT high-quality clusters, proTRAC, and PILFER in humans, mice, and flies (fly annotations only available from piRNAdb) using small RNA-seq datasets from ovaries, testes, and somatic tissues.

|  |  | piRNAdb |  |  | Girard et al. 2006 |  |  |
| --- | --- | --- | --- | --- | --- | --- | --- |
|  |  | piRAT-HQ | proTRAC | PILFER | piRAT-HQ | proTRAC | PILFER |
| **Human** |  |  |  |  |  |  |  |
|  | **Testes** | 93.65% | 36.91% | 54.50% | 66.40% | 25.95% | 37.87% |
|  | **Ovaries** | 33.57% | 5.54% | 1.58% | 16.42% | 2.29% | 3.04% |
|  | **Somatic** | 0.00% | 0.00% | 0.00% | 0.00% | 0.00% | 0.00% |
| **Mouse** |  |  |  |  |  |  |  |
|  | **Testes** | 6.85% | 4.64% | 22.88% | 82.97% | 75.60% | 80.39% |
|  | **Ovaries** | 3.17% | 0.00% | 4.94% | 5.61% | 0.00% | 6.77% |
|  | **Somatic** | 0.00% | 0.00% | 2.04% | 0.00% | 0.00% | 0.96% |
| **Fly** |  |  |  |  |  |  |  |
|  | **Testes** | 62.90% | 0.00% | 6.90% |  |  |  |
|  | **Ovaries** | 81.96% | 4.30% | 95.80% |  |  |  |
|  | **Somatic** | 0.22% | 0.00% | 3.61% |  |  |  |

### 3.4 Computing performance

While running the benchmarks, we monitored the running time and the maximum RAM usage in the same machine using a single thread, as well as 26 threads in piRAT (**Supplementary Table S4**). Running with a single thread piRAT takes the longest.

However, piRAT running time can be significantly shortened using multiple threads. For example, for the four datasets of mouse testes for piRAT it took ∼29h, proTRAC ∼3h, and PILFER ∼19h, but piRAT running time was decreased to ∼2h using 26 threads.

### 3.5 Overall performance

Based on our comparison, piRAT emerges as the most complete tool in terms of functionalities and supporting information that it provides. The benchmarking shows that it consistently delivers higher-quality annotations of piRNA clusters and ping-pong sites compared to alternative tools. Because it is easy to use, doesn’t need prior annotations or special pre-processing, and provides high-quality annotations and reports, we believe it is an excellent choice for any species, but especially for non-model organisms.

## Acknowledgments

We thank all the lab members who have been involved in technical discussions at different stages of the development.

## Author contributions

DR wrote the piRAT’s code, performed analysis, and prepared the figures. GY conceived and led the project and wrote the manuscript. All authors approved the final version of the manuscript.

## References

Aravin AA, Hannon GJ, Brennecke J. The Piwi-piRNA Pathway Provides an Adaptive Defense in the Transposon Arms Race. Science 2007;318:761–4.

Aravin AA, Naumova NM, Tulin AV et al. Double-stranded RNA-mediated silencing of genomic tandem repeats and transposable elements in the D. melanogaster germline. Current Biology 2001;11:1017–27.

Brennecke J, Aravin AA, Stark A et al. Discrete Small RNA-Generating Loci as Master Regulators of Transposon Activity in Drosophila. Cell 2007;128:1089–103.

Carmell MA, Xuan Z, Zhang MQ et al. The Argonaute family: tentacles that reach into RNAi, developmental control, stem cell maintenance, and tumorigenesis. Genes Dev 2002;16:2733–42.

Chen S, Ben S, Xin J et al. The biogenesis and biological function of PIWI-interacting RNA in cancer. Journal of Hematology & Oncology 2021;14:93.

Czech B, Hannon GJ. One Loop to Rule Them All: The Ping-Pong Cycle and piRNA-Guided Silencing. Trends in Biochemical Sciences 2016;41:324–37.

Ester M, Kriegel H-P, Sander J et al. A Density-Based Algorithm for Discovering Clustersin Large Spatial Databases with Noise. Proceedings of 2nd International Conference on Knowledge Discovery and Data Mining 1996:226–31.

Gainetdinov I, Colpan C, Arif A et al. A Single Mechanism of Biogenesis, Initiated and Directed by PIWI Proteins, Explains piRNA Production in Most Animals. Molecular Cell 2018;71:775-790.e5.

Girard A, Sachidanandam R, Hannon GJ et al. A germline-specific class of small RNAs binds mammalian Piwi proteins. Nature 2006;442:199–202.

Gunawardane LS, Saito K, Nishida KM et al. A Slicer-Mediated Mechanism for Repeat-Associated siRNA 5’ End Formation in Drosophila. Science 2007;315:1587–90.

Houwing S, Berezikov E, Ketting RF. Zili is required for germ cell differentiation and meiosis in zebrafish. The EMBO Journal 2008;27:2702–11.

Jung I, Park JC, Kim S. PiClust: A density based piRNA clustering algorithm. Computational Biology and Chemistry 2014;50:60–7.

Kawaoka S, Arai Y, Kadota K et al. Zygotic amplification of secondary piRNAs during silkworm embryogenesis. RNA 2011;17:1401–7.

Kawaoka S, Hayashi N, Katsuma S et al. Bombyx small RNAs: Genomic defense system against transposons in the silkworm, Bombyx mori. Insect Biochemistry and Molecular Biology 2008;38:1058–65.

Kiuchi T, Koga H, Kawamoto M et al. A single female-specific piRNA is the primary determiner of sex in the silkworm. Nature 2014;509:633–6.

Lau NC, Seto AG, Kim J et al. Characterization of the piRNA complex from rat testes. Science 2006, DOI: 10.1126/science.1130164.

Lewis SH, Quarles KA, Yang Y et al. Pan-arthropod analysis reveals somatic piRNAs as an ancestral defence against transposable elements. Nature Ecology & Evolution 2018;2:174–81.

Liu Y, Dou M, Song X et al. The emerging role of the piRNA/piwi complex in cancer. Molecular Cancer 2019;18:123.

Llonga N, Ylla G, Bau J et al. Diversity of piRNA expression patterns during the ontogeny of the German cockroach. Journal of Experimental Zoology Part B: Molecular and Developmental Evolution 2018;330:288–95.

Mito T, Ishimaru Y, Watanabe T et al. Cricket: The third domesticated insect. Current Topics in Developmental Biology. Vol 147. Academic Press, 2022, 291–306.

Montañés JC, Rojano C, Ylla G et al. siRNA enrichment in Argonaute 2-depleted Blattella germanica. Biochimica et Biophysica Acta - Gene Regulatory Mechanisms 2021;1864:194704.

Nakamura T, Ylla G, Extavour CG. Genomics and genome editing techniques of crickets, an emerging model insect for biology and food science. Current Opinion in Insect Science 2022;50:100881.

Piuco R, Galante PAF. piRNAdb: A piwi-interacting RNA database. 2021:2021.09.21.461238.

Raabe CA, Tang T-H, Brosius J et al. Biases in small RNA deep sequencing data. Nucleic Acids Research 2014;42:1414–26.

Ramat A, Simonelig M. Functions of PIWI Proteins in Gene Regulation: New Arrows Added to the piRNA Quiver. Trends in Genetics 2021;37:188–200.

Ray R, Pandey P. piRNA analysis framework from small RNA-Seq data by a novel cluster prediction tool - PILFER. Genomics 2018;110:355–65.

Rosenkranz D, Zischler H. proTRAC - a software for probabilistic piRNA cluster detection, visualization and analysis. BMC Bioinformatics 2012;13:5.

Uhrig S, Klein H. PingPongPro: a tool for the detection of piRNA-mediated transposon-silencing in small RNA-Seq data. Bioinformatics 2019;35:335–6.

Weng W, Li H, Goel A. Piwi-interacting RNAs (piRNAs) and cancer: Emerging biological concepts and potential clinical implications. Biochimica et Biophysica Acta (BBA) - Reviews on Cancer 2019;1871:160–9.

Yamashita T, Komenda K, Miłodrowski R et al. Non-gonadal expression of piRNAs is widespread across Arthropoda. FEBS Letters 2025;599:3–18.

