## Supplementary Figures for "piRAT: piRNA Annotation Tool for annotating, analyzing, and visualizing piRNAs"

### Supplementary Figure S1

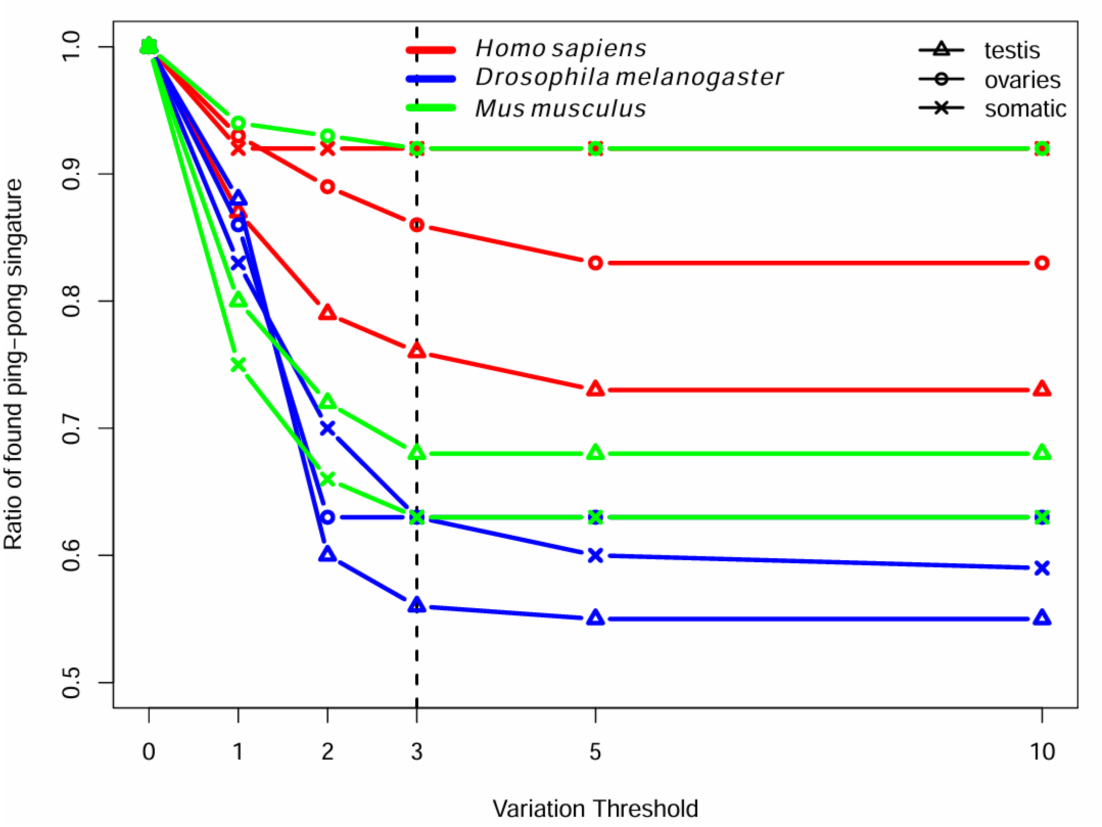


**Supplementary Figure S1:** Ratio of identified ping-pong signatures as a function of the variation threshold (number of different bases between reads mapped in the same locus). Beyond 3 nucleotide differences, the number of ping-pong signatures stabilizes.

**Supplementary Figure 2**

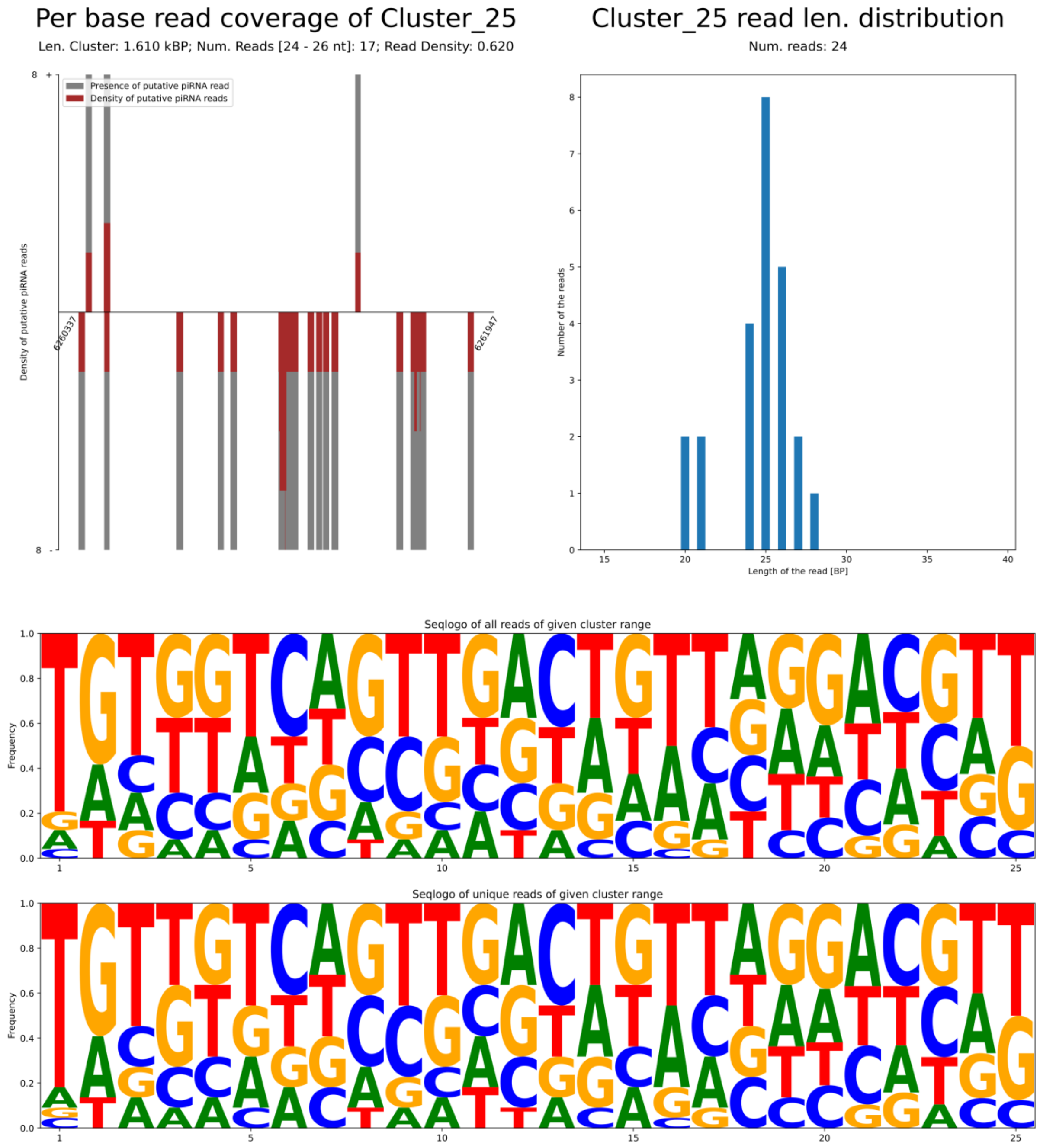

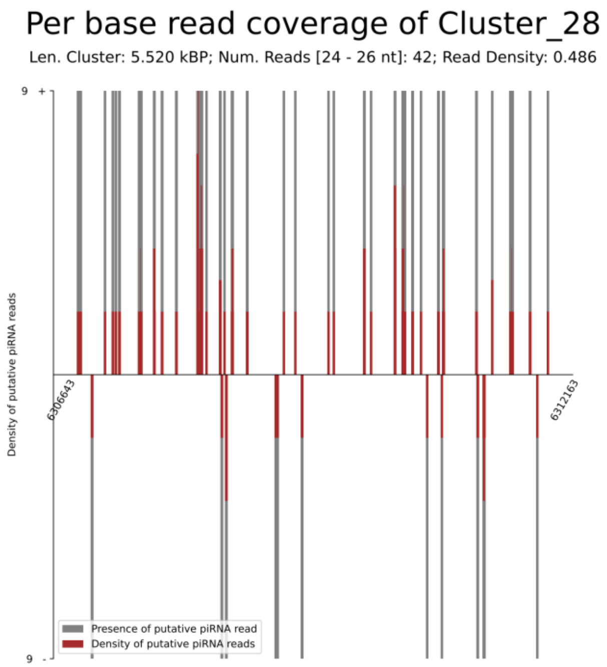
Per base read coverage of Cluster_28
Len. Cluster: 5.520 kBP; Num. Reads [24 - 26 nt]: 42; Read Density: 0.486
Density of putative piRNA reads
9 +
9 -
6306643
6312163
Presence of putative piRNA read
Density of putative piRNA reads

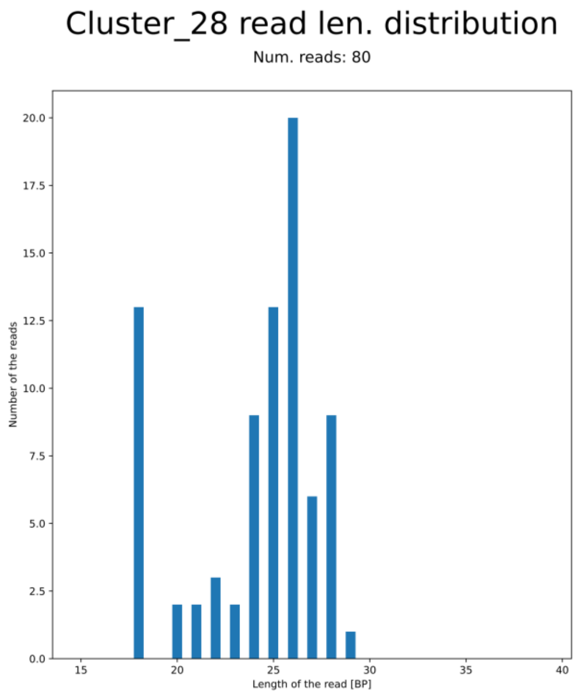
Cluster_28 read len. distribution
Num. reads: 80
Number of the reads
Length of the read [BP]
0.0
2.5
5.0
7.5
10.0
12.5
15.0
17.5
20.0
15
20
25
30
35
40

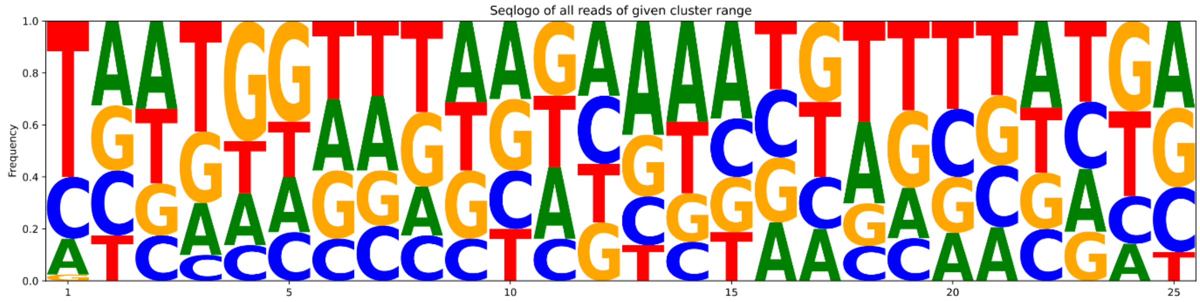
Seqlogo of all reads of given cluster range
Frequency
0.0
0.2
0.4
0.6
0.8
1.0
1
5
10
15
20
25

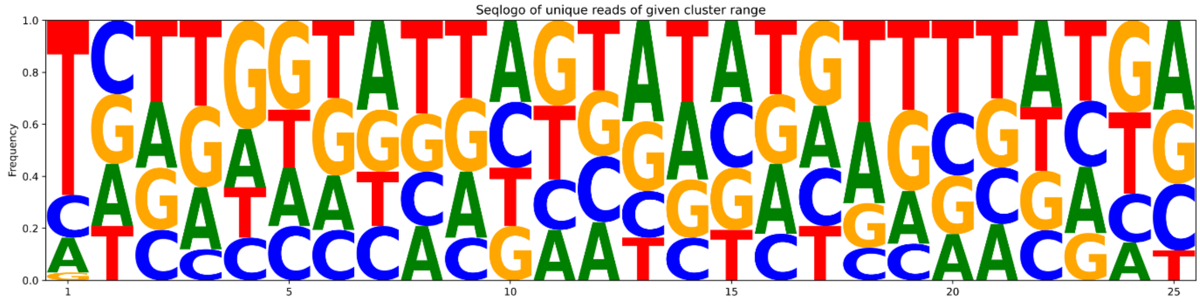
Seqlogo of unique reads of given cluster range
Frequency
0.0
0.2
0.4
0.6
0.8
1.0
1
5
10
15
20
25

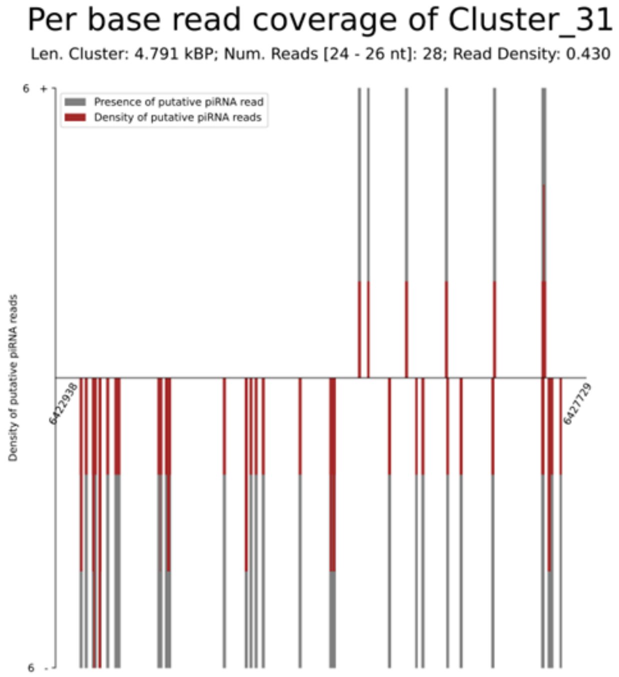
Per base read coverage of Cluster_31
Len. Cluster: 4.791 kBP; Num. Reads [24 - 26 nt]: 28; Read Density: 0.430
Presence of putative piRNA read
Density of putative piRNA reads
Density of putative piRNA reads
6 +
6 -
6422938
6427729

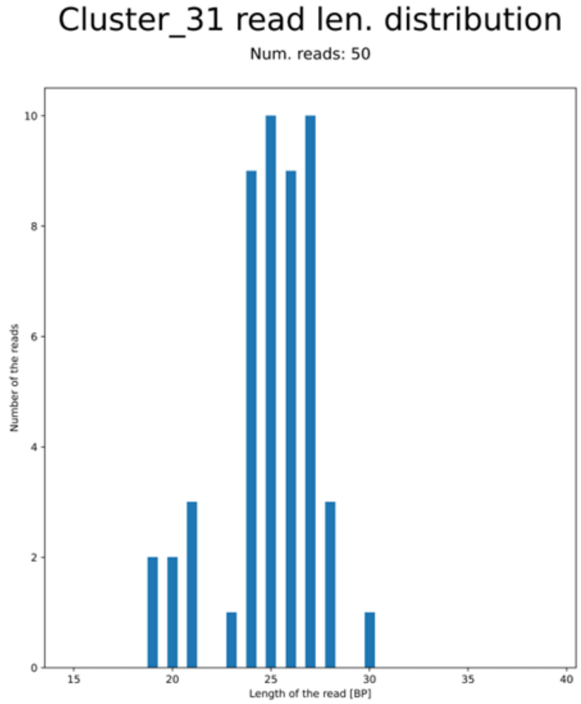
Cluster_31 read len. distribution
Num. reads: 50
Number of the reads
Length of the read [BP]

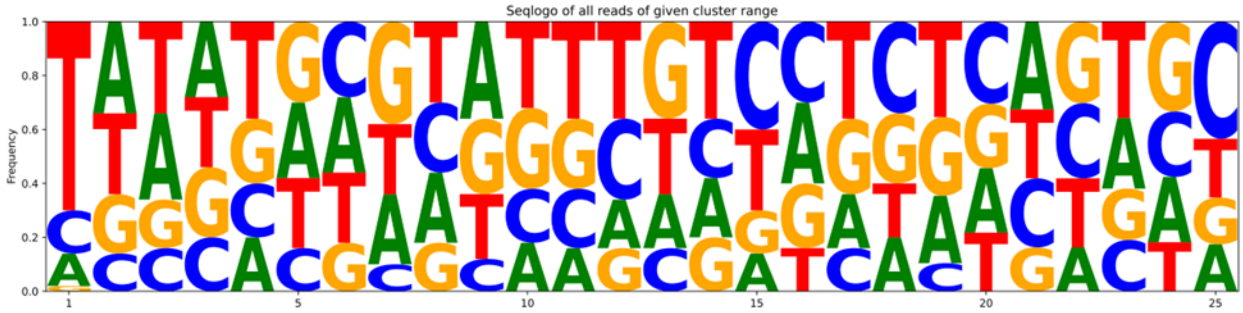
Seqlogo of all reads of given cluster range
Frequency

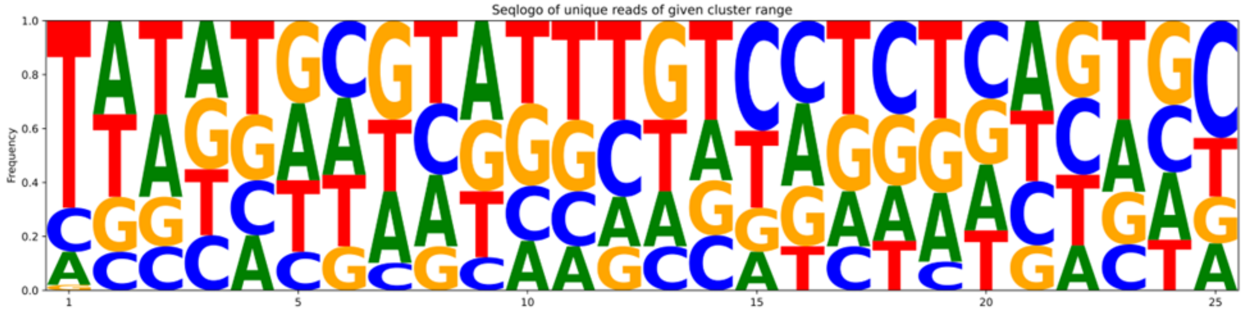
Seqlogo of unique reads of given cluster range
Frequency

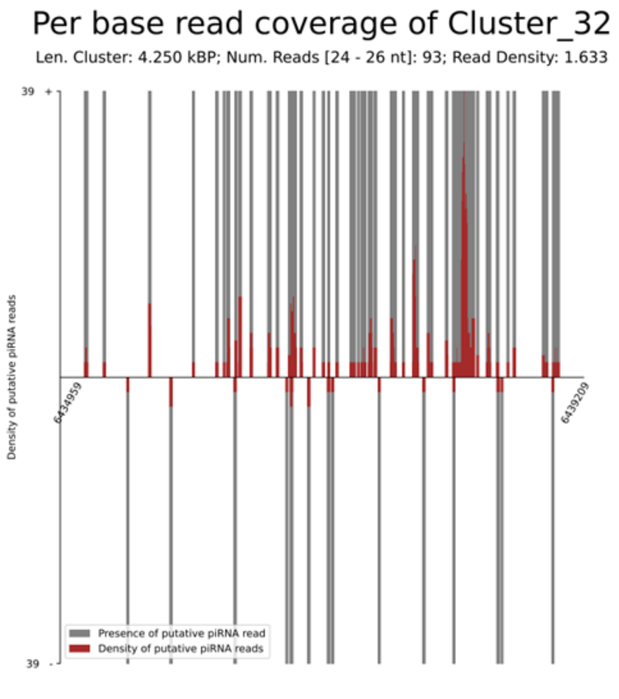
Per base read coverage of Cluster_32
Len. Cluster: 4.250 kBP; Num. Reads [24 - 26 nt]: 93; Read Density: 1.633
39 +
39 -
Density of putative piRNA reads
6434959
6439209
Presence of putative piRNA read
Density of putative piRNA reads

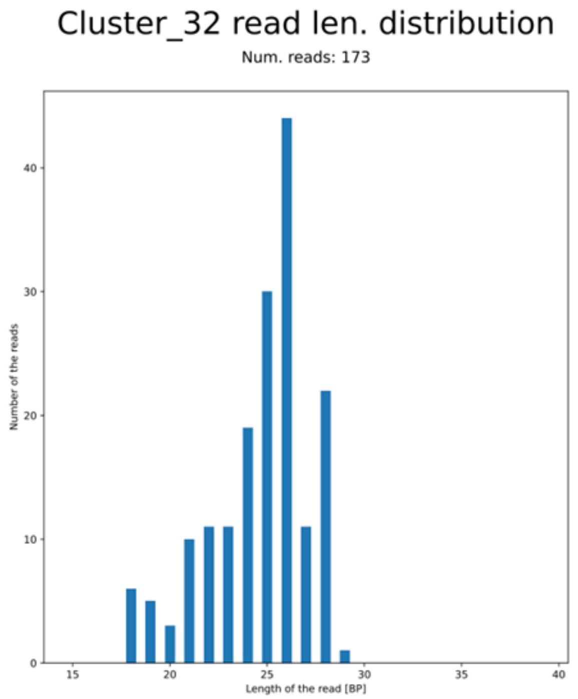
Cluster_32 read len. distribution
Num. reads: 173
Number of the reads
Length of the read [BP]

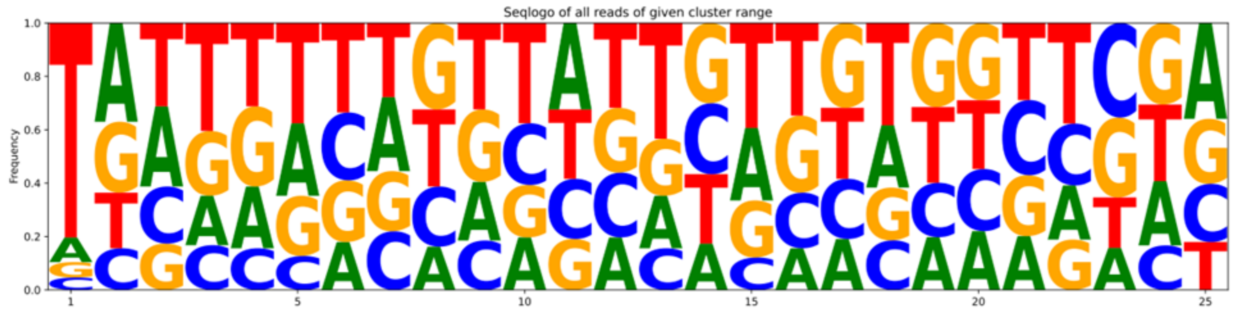
Seqlogo of all reads of given cluster range
Frequency

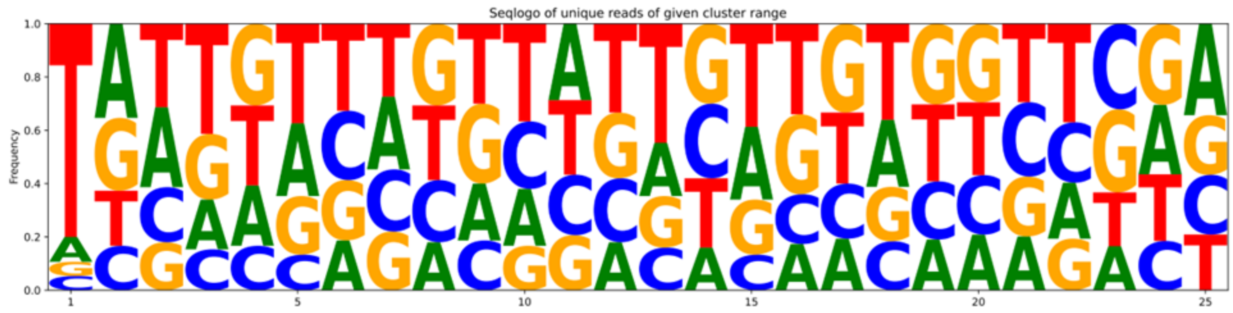
Seqlogo of unique reads of given cluster range
Frequency

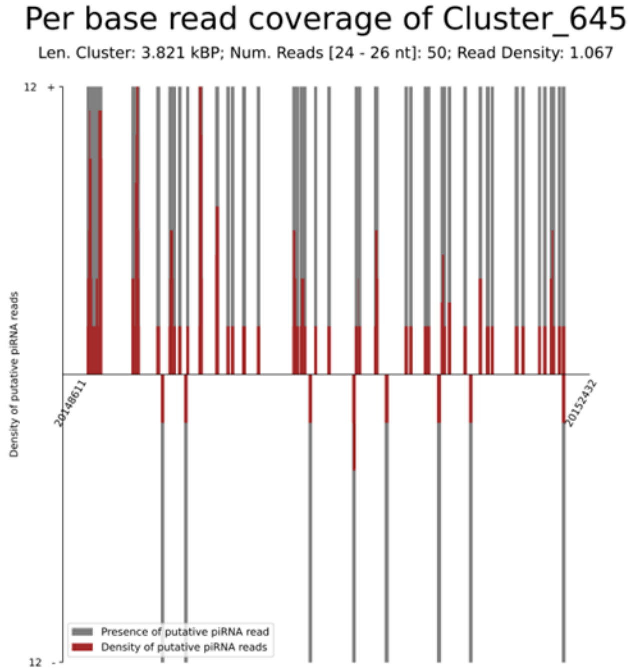
Per base read coverage of Cluster_645
Len. Cluster: 3.821 kBP; Num. Reads [24 - 26 nt]: 50; Read Density: 1.067
Density of putative piRNA reads
20148611
20152432
Presence of putative piRNA read
Density of putative piRNA reads

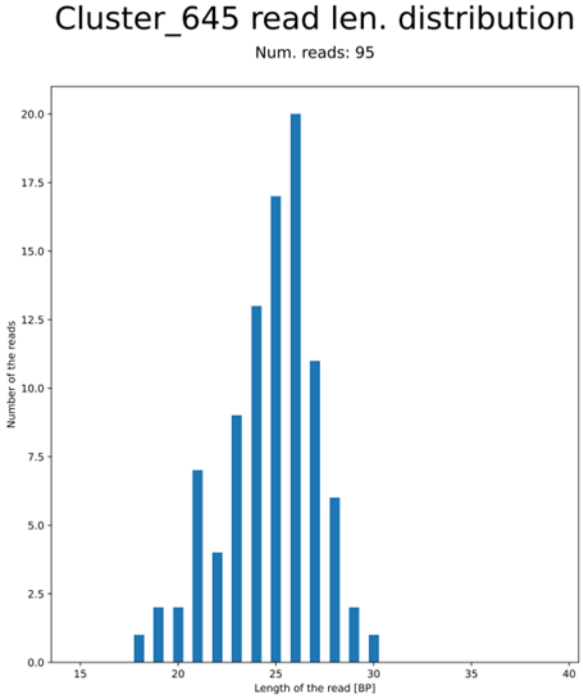
Cluster_645 read len. distribution
Num. reads: 95
Number of the reads
Length of the read [BP]

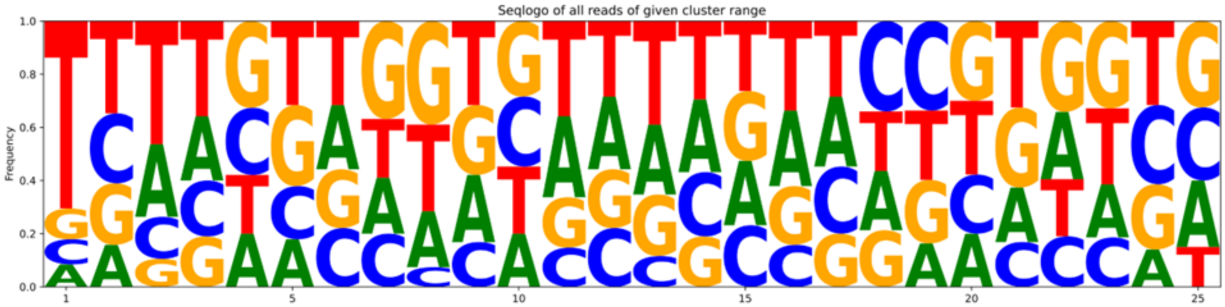
Seqlogo of all reads of given cluster range
Frequency

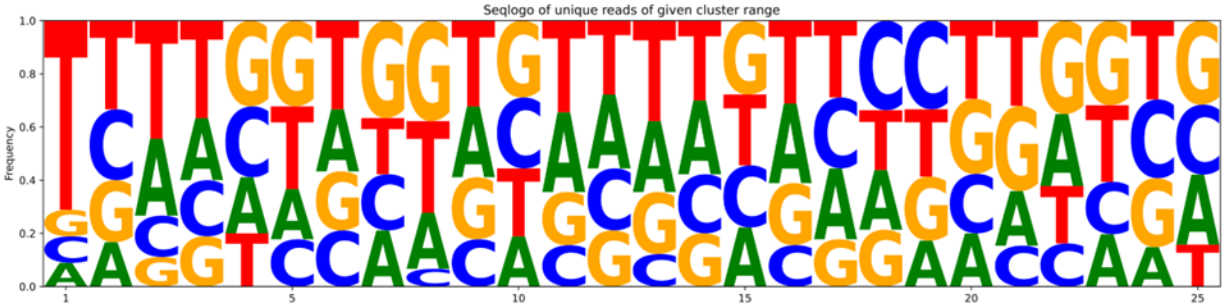
Seqlogo of unique reads of given cluster range
Frequency

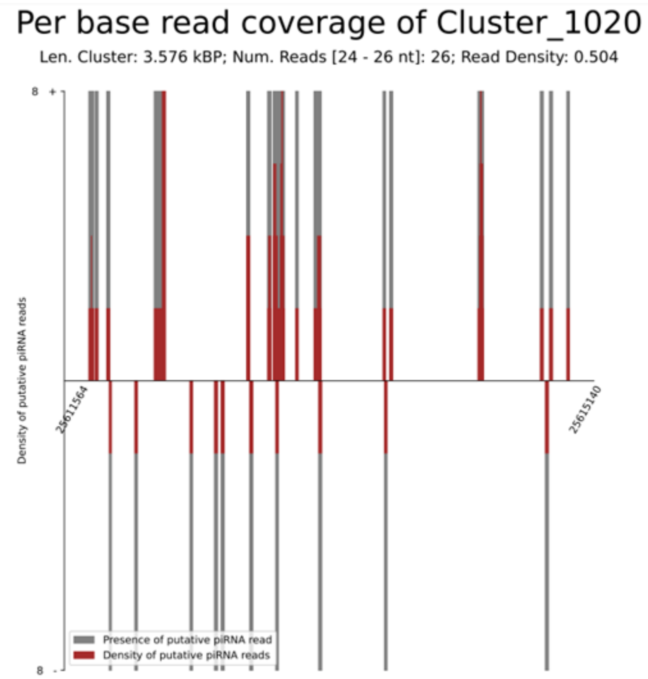
Per base read coverage of Cluster_1020
Len. Cluster: 3.576 kBP; Num. Reads [24 - 26 nt]: 26; Read Density: 0.504
Density of putative piRNA reads
25611564
25615140
Presence of putative piRNA read
Density of putative piRNA reads

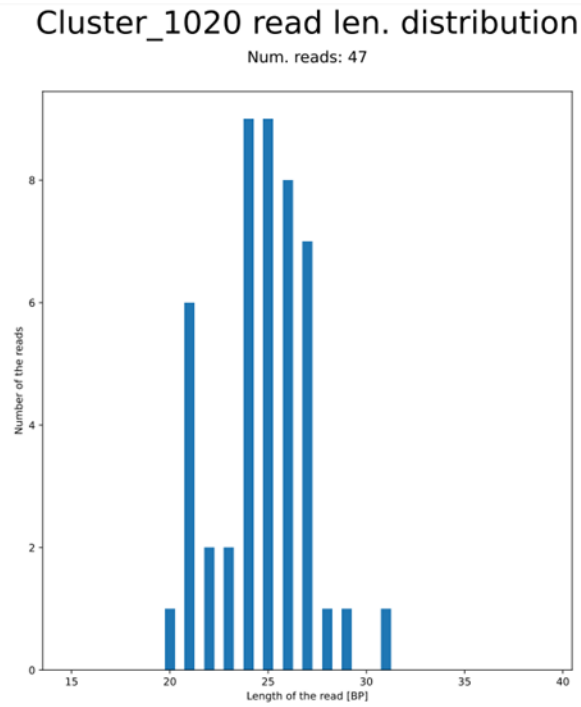
Cluster_1020 read len. distribution
Num. reads: 47
Number of the reads
Length of the read [BP]

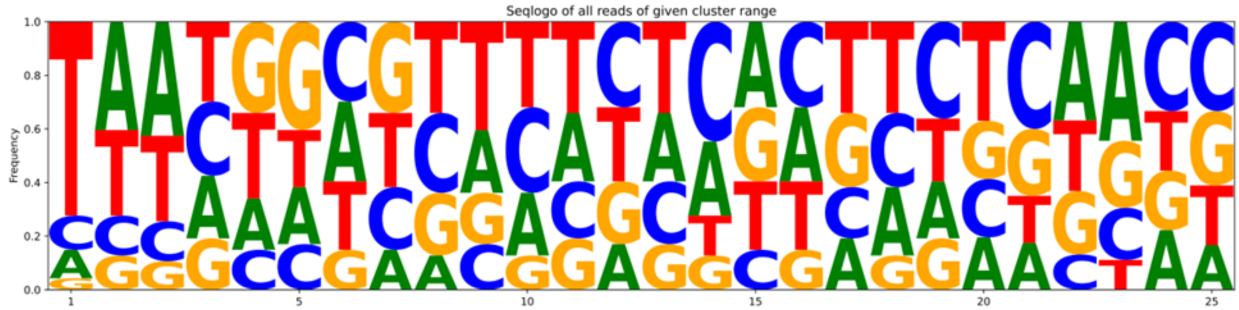
Seqlogo of all reads of given cluster range
Frequency

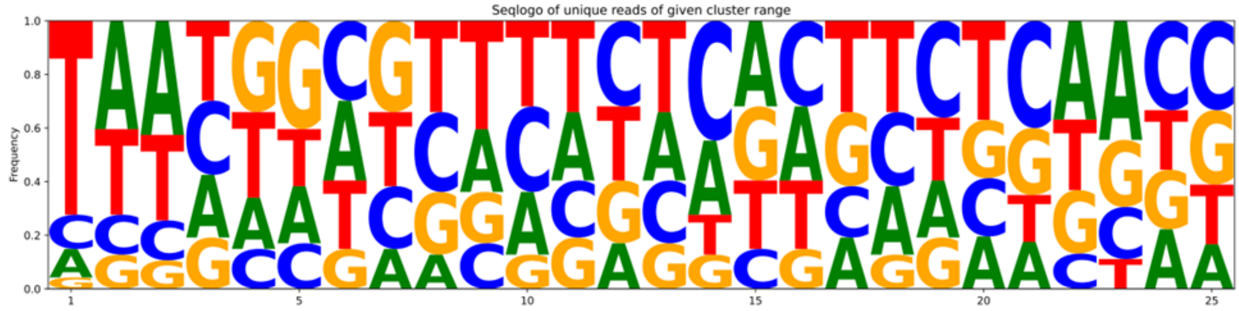
Seqlogo of unique reads of given cluster range
Frequency

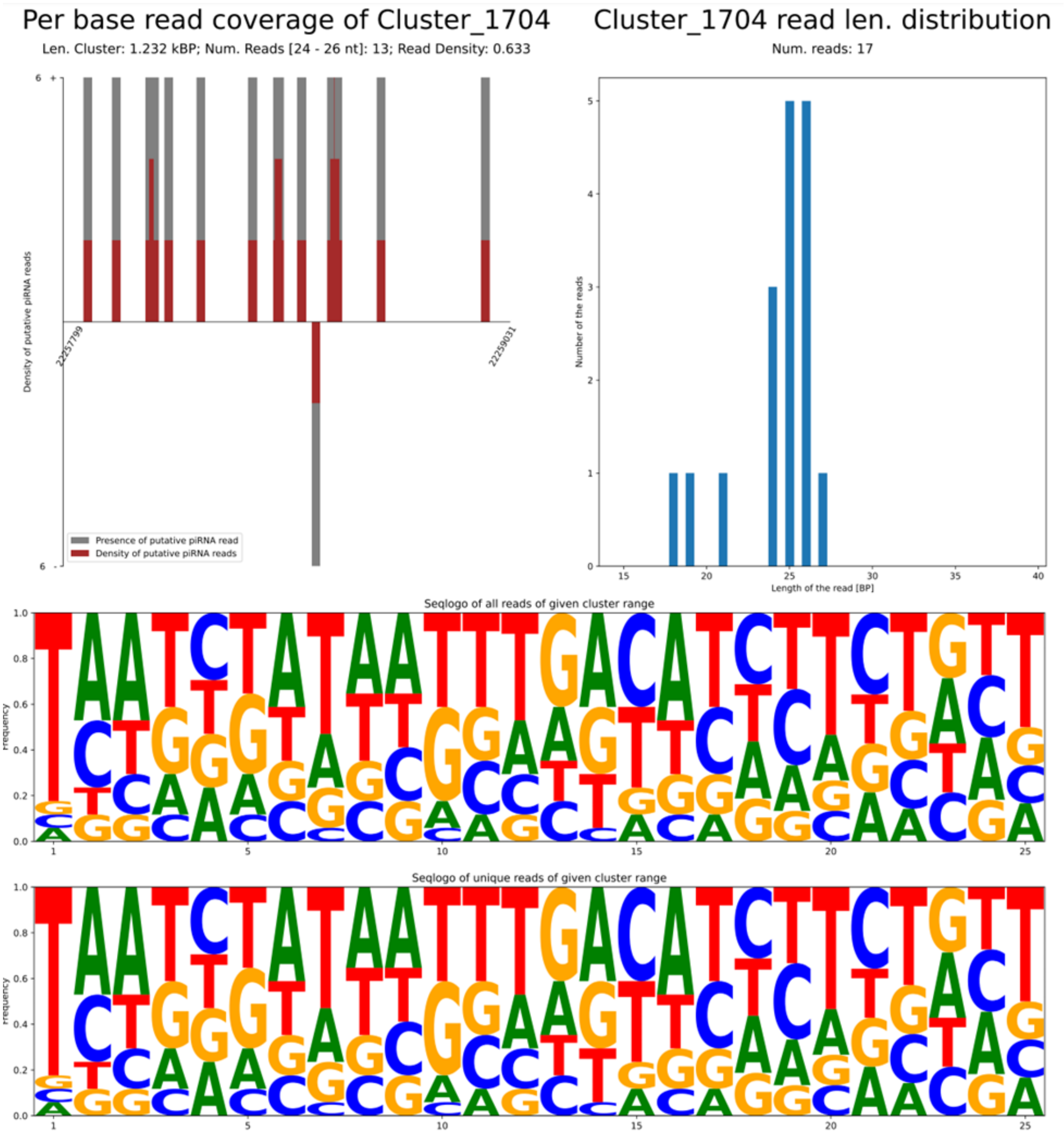

Per base read coverage of Cluster_1704
Len. Cluster: 1.232 kBP; Num. Reads [24 - 26 nt]: 13; Read Density: 0.633
Density of putative piRNA reads
22257799
22259031
Presence of putative piRNA read
Density of putative piRNA reads
Cluster_1704 read len. distribution
Num. reads: 17
Number of the reads
Length of the read [BP]
Seqlogo of all reads of given cluster range
Frequency
Seqlogo of unique reads of given cluster range
Frequency

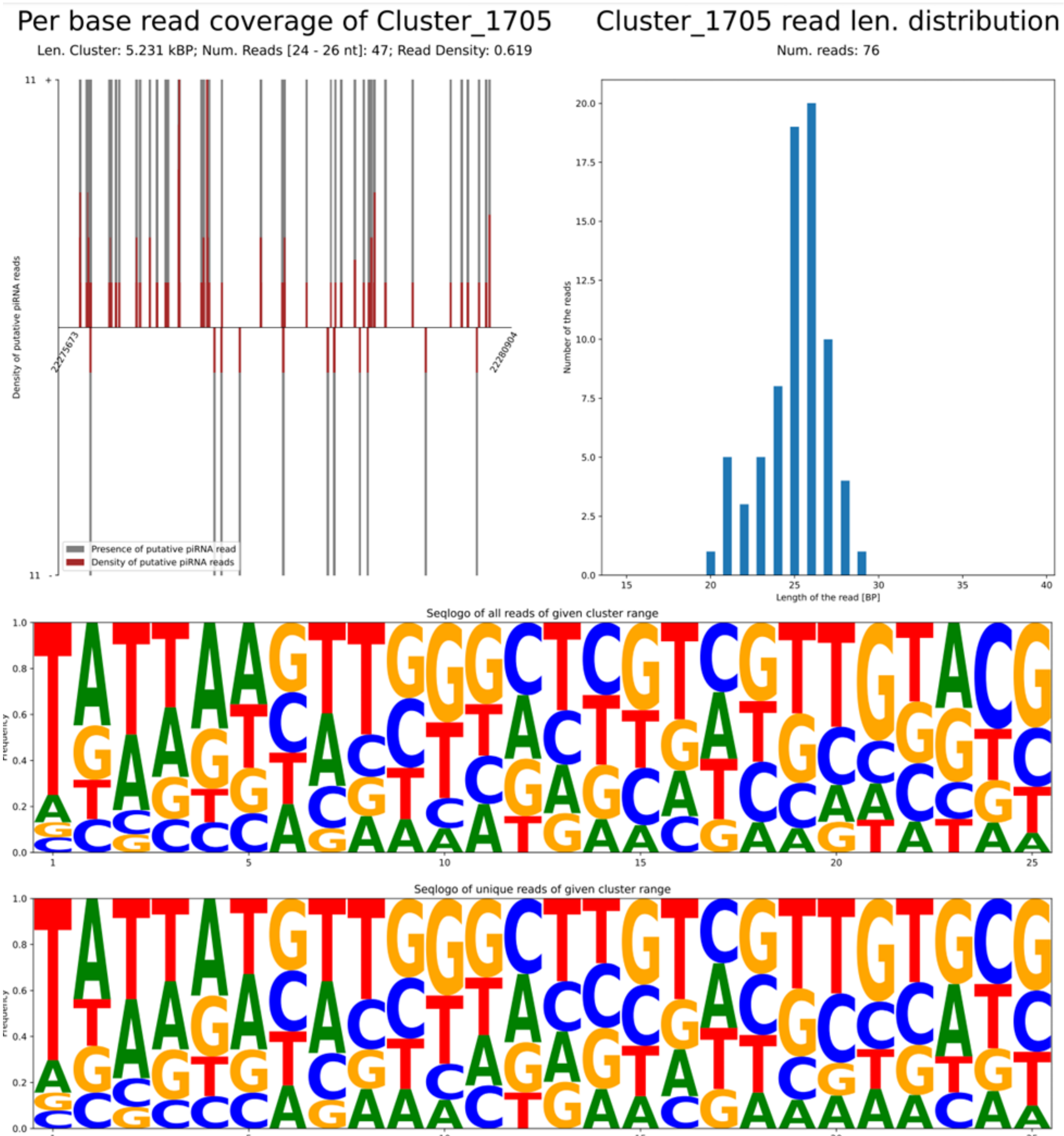

Per base read coverage of Cluster_1705
Len. Cluster: 5.231 kBP; Num. Reads [24 - 26 nt]: 47; Read Density: 0.619
Density of putative piRNA reads
22275673
22280904
Presence of putative piRNA read
Density of putative piRNA reads
Cluster_1705 read len. distribution
Num. reads: 76
Number of the reads
Length of the read [BP]
Seqlogo of all reads of given cluster range
Seqlogo of unique reads of given cluster range

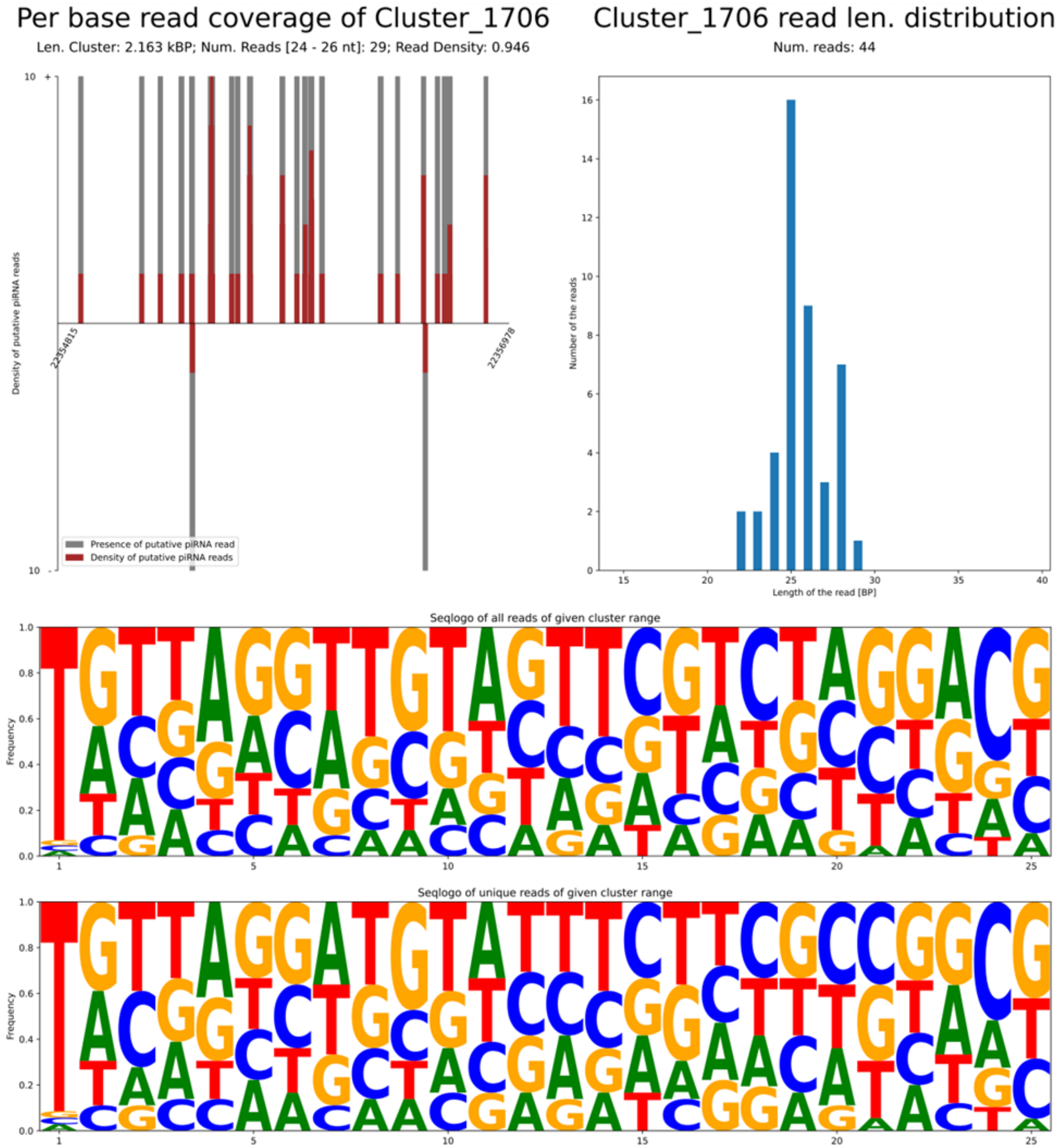
Per base read coverage of Cluster_1706
Len. Cluster: 2.163 kBP; Num. Reads [24 - 26 nt]: 29; Read Density: 0.946
Density of putative piRNA reads
22354815
22356978
Presence of putative piRNA read
Density of putative piRNA reads
Cluster_1706 read len. distribution
Num. reads: 44
Number of the reads
Length of the read [BP]
Seqlogo of all reads of given cluster range
Frequency
Seqlogo of unique reads of given cluster range
Frequency

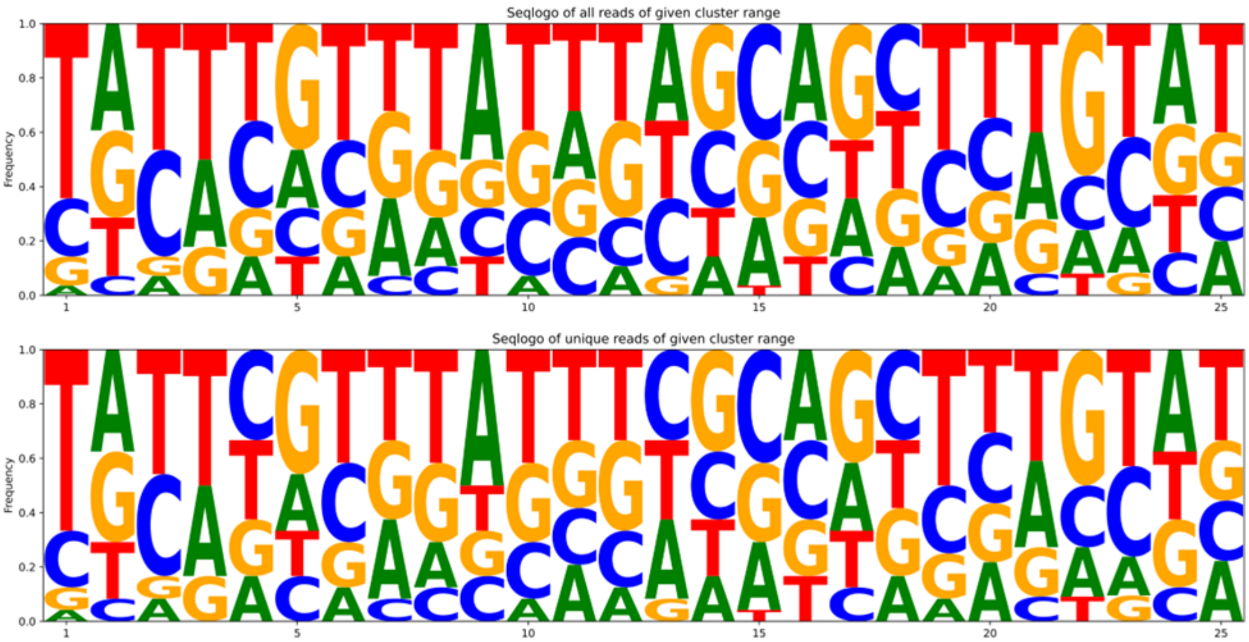


**Supplementary Figure S2**: Plots produced by piRAT supporting the annotation of the 9 somatic piRNA clusters in *Drosophila melanogaster*.
