## Supplementary Methods for "piRAT: piRNA Annotation Tool for annotating, analyzing, and visualizing piRNAs"

#### 1. piRNA Length Determination in piRAT

While the user can provide a range of piRNA length if it is known for the species they analyze, for non-model species, this information might not be available. When a piRNA length range is not provided, piRAT will predict it from small RNA-seq reads based on one of the two modes (depending on running mode):

##### a) Running mode : "Full Analysis" or "Ping-Pong Loci Identification"(flag: -m secondary)

When piRAT runs in the modes of full analysis (primary + secondary annotations) or only secondary, the piRNA range will be estimated based only on the length of reads exhibiting the characteristic ping-pong signature (10nt 5' to 5' overlap). For this, piRAT first processes each input BAM file individually to identify reads involved in ping-pong signatures and calculates z-scores for the abundance of each read length participating in these signatures. Then, for each specific length, these z-scores are averaged across all analyzed samples. The final piRNA range for the dataset is then defined to include all lengths where this average z-score surpasses the threshold of 0.5 (**Supplementary Methods Figure 1** ).

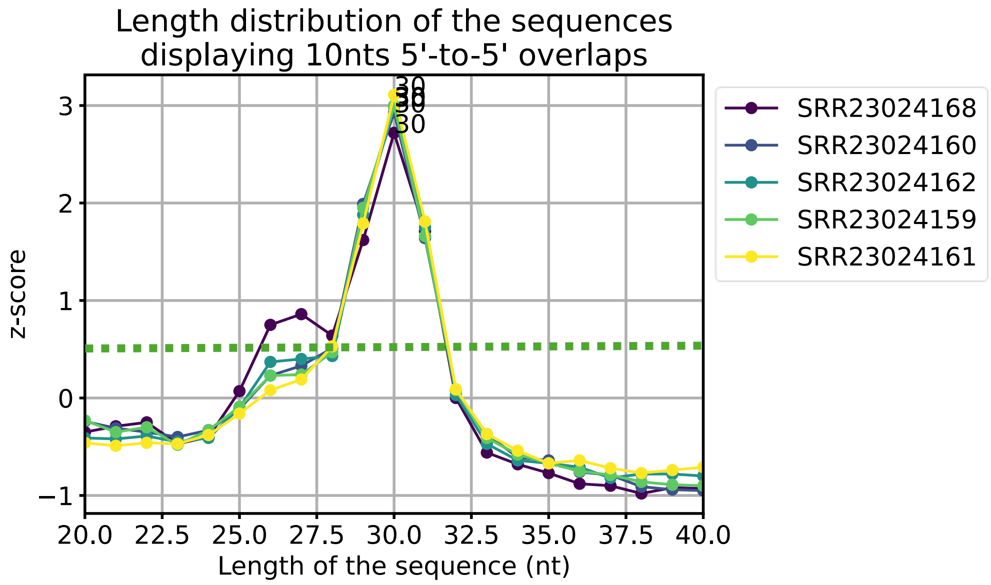

**Supplementary Methods Figure 1:** Z-score length distribution of reads displaying ping-pong signature (10 nt overlaps 5'-5'), demonstrating enrichment of specific piRNA lengths in the representative dataset. Peaks with Z-score > 0.5 represent lengths significantly enriched and thus selected as putative piRNA length range in the given species (in this case the defined range would be 27-31).

##### b) Running mode: "Primary piRNA Cluster Annotation" (flag: -m primary)

When piRAT runs only in "primary mode" (not performing ping-pong annotations) it will randomly select 100,000 mapped reads from each bam file. It then calculates the count distribution of sequence lengths and identifies the predominant length (highest peak) within the range of 26 to 32 nucleotides (nt). After the predominant piRNA length is defined, piRAT determines the range of piRNA lengths based on the weighted mean: piRAT assigns a weight of 1.0 to the count of the predominant length (the highest peak). Subsequently, other nucleotide lengths within the specified range (26-32) are ranked according to their respective read counts in descending order. A progressively decreasing weight is assigned to these ranked lengths: the length with the second-highest read count receives a weight of 0.9, the one with the third-highest count receives 0.8, and so forth. These weights are then utilized in the calculation of the weighted mean. The identified length distributions are illustrated in **Supplementary Methods Figure 2**, showing distinct peaks in the newly defined length range.

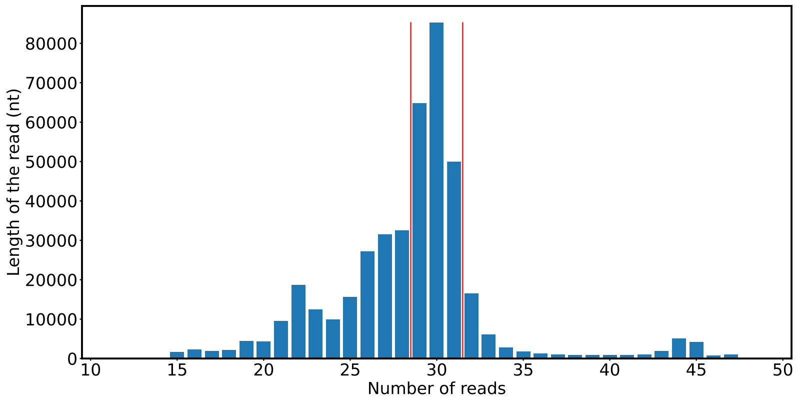

**Supplementary Methods Figure 2:** Read the length distribution of the 100,000 random reads used to estimate the piRNA length range when piRAT runs without computing pin-pong signatures (“Primary piRNA Cluster Annotation” mode). The bar at 30nts corresponds to the predominant piRNA length detected in this sample, and the length between the red vertical bars (29-31) indicates the predicted piRNA length range in thi given dataset.

### 2. Annotation of primary piRNA clusters

The annotation of primary piRNA clusters with piRAT, based on a modified version of the DBSCAN algorithm (Ester *et al.* 1996), involves five main steps: (a) purging reads, (b) identifying optimal clustering parameters, (c) clustering, (d) grouping and classification of clusters, and (e) quality analysis of annotated clusters

#### a) Purging the reads

After loading the reads from each file, piRAT purges them to reduce noise and ensure accuracy in subsequent steps, as described in **Section 2.1** of the main article.

#### b) Finding the optimal clustering parameters

There are two parameters in the DBSCAN algorithm that piRAT optimizes automatically. These are Eps and k:

- Eps: Maximum allowed gap (distance) between neighboring reads within a cluster.
- k: Minimum number of reads required within distance Eps to define a valid cluster core region.

To determine these values, piRAT calculates the distance for each read in the dataset (mapped to the same chromosome and strand) to its k-th (for k: {4, 6,8, 10} nearest neighboring read. (**Supplementary Methods Figure 3**)

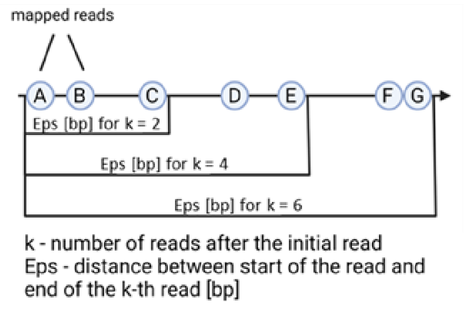

**Supplementary Methods Figure 3:** Scheme of method of calculating distances between k-reads.

From the calculated k-th nearest neighbor distances for a given k, piRAT initially keeps only those distances less than or equal to 10,000 nt. This value was chosen because biologically relevant piRNA clusters, while potentially long, are generally characterized by reads that are relatively close to each other, and it is uncommon for such clusters to span true internal gaps larger than 10 ,000 nt. Filtered k-th neighbor distances are then sorted in descending order.

To identify an optimal Eps, piRAT analyzes this sorted distribution to find a 'knee' or 'elbow' – a point where the distances begin to stop decreasing and plateaus (**Supplementary Methods Figure 4A**). This 'knee' typically distinguishes denser regions (inter-cluster distances) from sparser regions. To locate this point, piRAT divides the sorted distances into 20 segments and calculates the rate of change (slope) of distances between these segments (Supplementary Methods Figure 4B).

The Eps value for given k is then automatically selected as the distance value corresponding to the point just before this plateau. To select the most appropriate k from the set {4, 6, 8, 10} (and its corresponding Eps), piRAT further evaluates the characteristics of the plateau (in the same matter as above – the plateau region is divided into 20 parts, and its slope is calculated). The condition for a 'good' k is met if at least 50% of the segments constituting this plateau demonstrate a rate of change in slope that is greater than -1. Such a profile indicates that k-distances within the majority of the plateau are stable and not significantly decreasing, confirming a consistent, high-density region suitable for clustering with the current k.

Conversely, if less than 50% of the plateau region meets this flatness criterion, it suggests that a stable, uniform density has not been achieved for that k – it shows piRAT that there is an insufficient number of reads in the dataset for the given k. If multiple k successfully meet the "sufficient flat plateau" criterion, piRAT choses the highest k that meets this criteria. Varying parameters (Eps, k) influence cluster sensitivity and specificity. Users can optimize these settings based on experimental context or data quality, balancing the number of clusters detected and their confidence.

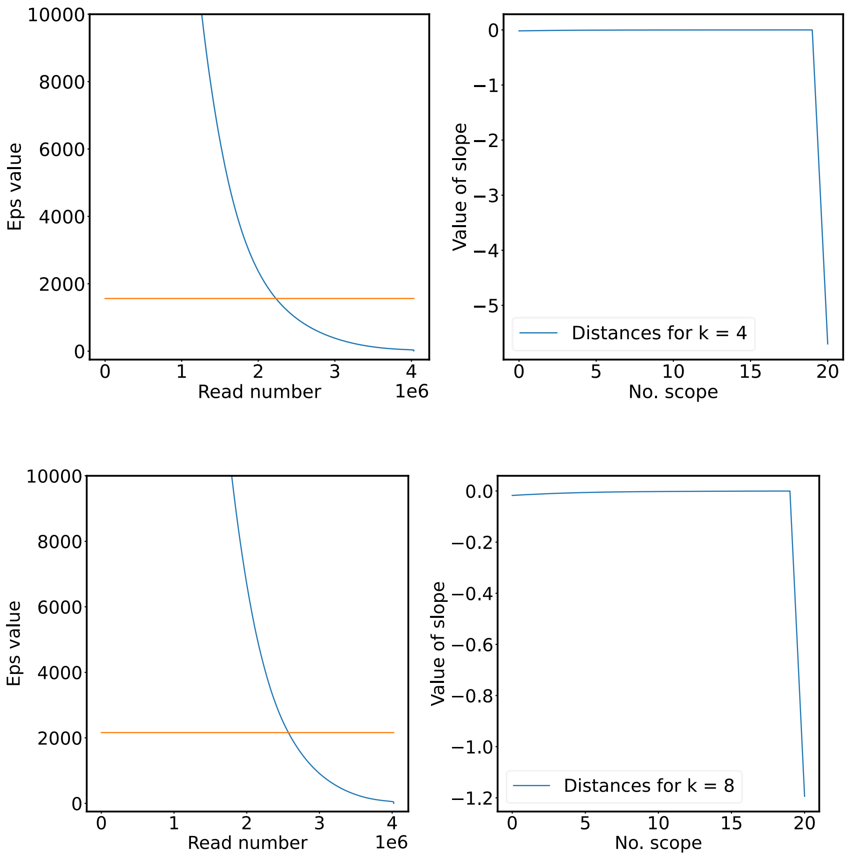

**Supplementary Methods Figure 4:** Automatic selection of DBSCAN's k and Eps parameters, illustrated with example for k=4 (optimal) and k=8 (suboptimal). Left – distribution of k-th nearest neighbor distances sorted in descending order. Calculated Eps is shown with an orange line. Right – rate of change of slope calculated for segments from the k-th nearest neighbor distances data. Both of the k values indicate a 'knee' in the same segment of the data, but the increase in k from 4 to 8, shows that for k=8, there is no sufficient plateau region within identified < Eps region.

#### c) Clustering

Due to sequencing artifacts, mapping ambiguities, or other sample preparation-related issues, sequencing data can exhibit variations in read coverage across a locus. This may lead to minor gaps within what is biologically a single, continuous piRNA-producing region. These 'minor gaps' can manifest as short segments with temporarily reduced read density, where an individual read might be spatially close (within an Eps distance) to an established cluster core, but does not itself meet the density requirement (MinReads) to be a core point. When encountered by a traditional density-based clustering algorithm – DBSCAN, these minor gaps can cause the premature fragmentation of a single, biologically relevant piRNA cluster into multiple smaller ones.

To address this, we modified the standard DBSCAN algorithm to specifically accommodate small discontinuities without prematurely terminating cluster growth, preserving biologically relevant clusters. The modifications ensure that, even with incomplete sequencing or uneven coverage, clusters are not fragmented by small gaps in the cluster coverage, as illustrated in **Supplementary Methods Figure 5**.

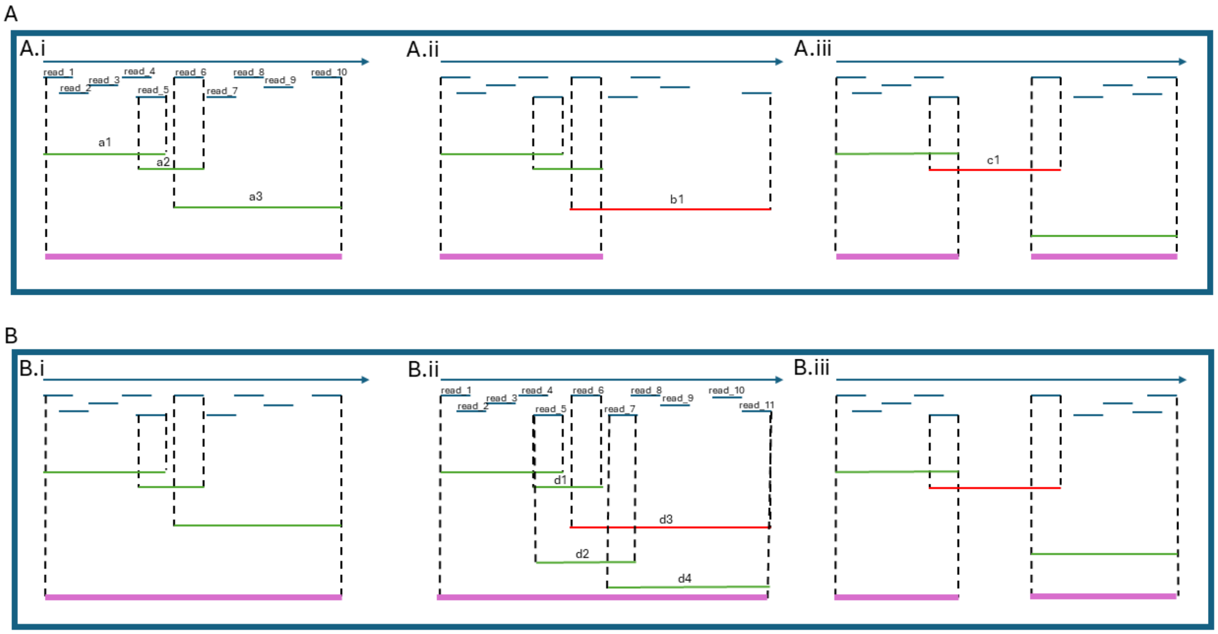

**Supplementary Methods Figure 5:** Comparison between the traditional DBSCAN algorithm (A) and our modified implementation ( B). In both versions, green lines indicate distances less than the Eps threshold (Eps < 1000), red lines indicate distances greater than the threshold (Eps > 1000), and magenta lines represent defined clusters. The parameters used for this example are MinReads = 4 and Eps = 1000.
(A) In the original DBSCAN algorithm, the process begins by evaluating the distance between the start of the first read (read_1) and the end of the read that is MinReads away (read_5). A new cluster core is defined if this distance is within the Eps parameter (A.i.a1). The algorithm then checks the next read (read_6) for continuity with the existing cluster. If the distance between the last read of the core (read_5) and the end of read_6 is within Eps (A.i.a2), the algorithm proceeds by checking the MinReads condition (A.i.a3). If this distance lies within Eps distance, the cluster core is expanded.
(A.ii) If the MinReads condition is not met for the next read outside of cluster core (A.ii.b1), this read (read_6) is marked as the border read, and the cluster expansion stops.
(A.iii) If a read lies outside the Eps distance (A.iii.c1), the expansion is stopped, and the previously defined cluster remains.
In our modified DBSCAN implementation, Panels (B) correspond to cases A.i, A.iii) from the original algorithm, with a key modification occurring in case (B.ii).
(B.i, B.iii) Cases in our modified algorithm behave identically to the original DBSCAN (A.i, A.iii), expanding the cluster when distances fall within the Eps threshold.
(B.ii) When the distance between the cluster core and the next read lies within Eps (B.ii.c1), but the MinReads condition is not met (B.ii.e3), the algorithm proceeds by evaluating the subsequent read (read_7). If both conditions — distance within Eps (B.ii.e)) and MinReads (B.ii.e4)) are satisfied, the cluster core is expanded.

The clustering process is performed individually per strand. The resulting clusters are saved in *clusters_out.gff*, with all associated reads stored in *Clusters_reads.gff*. Purged associated reads are stored in *potential_pirna_reads.gff*.

#### *i. Uni-strand, dual-strand, and bi-directional clusters.*

Because piRAT clustering algorithm runs in a strand-specific manner, the next step is to determine whether some clusters in close proximity but on opposite strands might need to be joint in a bidirectional or dual-strand cluster as follows:

- If two clusters in opposite strands overlap by at least half the length of the smaller cluster, or if the smaller cluster is entirely contained within the larger one, they are merged into a **dual-strand cluster**.
- If more than two clusters meet the overlap conditions, all of them are merged into a single **dual-strand cluster**.
- If two clusters are in opposite strands within an Eps distance of each other and do not fulfill the dual-strand criteria, they are merged into a **bi-directional cluster**.
- Clusters not meeting any of these criteria are classified as **uni-strand clusters**.

##### *ii. Quality analysis of annotated clusters*

Annotated clusters are classified in two categories, low-quality and high-quality clusters, based on piRNA biogenesis information, such as the frequency of thymine (T) as the first nucleotide in the piRNA reads and the length of the reads mapped within an annotated cluster.

High-quality clusters are those that:

- Have at least 50% of reads within the defined piRNA length range, and
- The frequency of thymine as the first nucleotide in all the reads belonging to the cluster is at least 50%.

High-quality clusters are stored in *high_quality_clusters_out.gff*, other clusters are stored in *low_quality_clusters_out.gff*.

### 3. Identifying the ping-pong loci

The identification of ping-pong loci consists of three main steps: (1) selection of representative reads, (2) detection of overlapping reads, and (3) identifying 10 nt overlap at the 5' ends.

- Selection of representative reads: After loading the reads from each file, piRAT detects the representative read per loci to reduce noise and ensure accuracy in subsequent steps. Further details on this step process can be found in Section 2.1 of the main manuscript.
- Measuring 5'-end overlaps: piRAT then calculated the length of any overlaps at the 5' ends and marks those reads that show potential 10-nt overlaps
- Filtering for 10-nt overlaps: From the flagged reads, piRAT select only those pairs with a confirmed 10-nt overlap (5' to 5')

The final list of identified ping-pong reads is saved to *ping_pong_reads.gff*
