## Supplementary Tables for "piRAT: piRNA Annotation Tool for annotating, analyzing, and visualizing piRNAs"

**Supplementary Table S1**: Publicly available small RNA-seq datasets used for benchmarking from human, mouse, and fly, from gonadal and non-gonadal tissues.

| | | | |
|---|---|---|---|
| **Human** | | | |
| | **Testis** | | **Sample accession code** |
| | | | ERR4846433 |
| | | | ERR4846432 |
| | **Ovaries** | | |
| | | | SRR1755253 |
| | | | SRR1755254 |
| | **Somatic** | muscle | |
| | | | SRR21813493 |
| | | | SRR21813494 |
| **Mouse** | | | |
| | **Testis** | | |
| | | | SRR23024159 |
| | | | SRR23024160 |
| | | | SRR23024161 |
| | | | SRR23024162 |
| | **Ovaries** | | |
| | | | SRR8494568 |
| | | | SRR8494567 |
| | **Somatic** | striatum | |
| | | | SRR13277520 |
| | | | SRR13277522 |
| | | | SRR13277523 |
| **Fly** | | | |
| | **Testis** | | |
| | | | SRR12213370 |
| | | | SRR22857968 |
| | **Ovaries** | | |
| | | | SRR7945626 |
| | | | SRR23073407 |
| | **Somatic** | head-thorax | |
| | | | SRR8539594 |
| | | | SRR7633524 |
| | | | SRR20830387 |

**Supplementary Table S2**: Percentage of small RNA-seq reads of the piRNA length (26-32 nts) and starting with T, mapped within the clusters among all the mapped reads in the clusters. For each species and tissue, using the clusters annotated by piRAT high-quality, proTRAC, and PILFER. NAs appear when no clusters are found in the given category.

| | | piRAT -high quality | proTRAC | PILFER |
|---|---|---|---|---|
| Human | | | | |
| | Testes | 65.91% | 68.00% | 16.29% |
| | Ovaries | 62.48% | 69.07% | 4.91% |
| | Somatic | NA | NA | 0.03% |
| Mouse | | | | |
| | Testes | 54.42% | 54.59% | 47.78% |
| | Ovaries | 48.35% | 51.50% | 0.97% |
| | Somatic | NA | NA | 0.21% |
| Fly | | | | |
| | Testes | 59.11% | 69.71% | 31.76% |
| | Ovaries | 49.51% | 52.39% | 49.00% |
| | Somatic | 43.79% | NA | 4.88% |

**Supplementary Table S3**: Percentage of small RNA-seq reads, of the piRNA length (26-32 nts) and starting with T, mapped within the annotated clusters among all the RNA-seq reads of the same characteristics mapped in the genome. For each species and tissue, using the clusters annotated by piRAT high-quality, proTRAC, and PILFER. NAs appear when no clusters are found in the given category.

| | | piRAT -high quality | proTRAC | PILFER |
|---|---|---|---|---|
| Human | | | | |
| | Testes | 90.00% | 10.54% | 12.42% |
| | Ovaries | 38.60% | 1.91% | 4.36% |
| | Somatic | NA | NA | 0.02% |
| Mouse | | | | |
| | Testes | 93.04% | 53.12% | 56.53% |
| | Ovaries | 35.85% | 0.24% | 0.51% |
| | Somatic | NA | NA | 0.04% |
| Fly | | | | |
| | Testes | 33.90% | 0.93% | 21.65% |
| | Ovaries | 59.71% | 1.03% | 28.38% |
| | Somatic | 0.00% | NA | 0.04% |

**Supplementary Table S4:** Computing time and maximum peak of RAM usage of piRAT, proTRAC, and PILFER while analyzing small RNA-seq data from testis, ovaries, and somatic tissue of human, fly, and mouse.

| | | **piRAT 26 threads** | | **piRAT 1 thread** | | **proTRAC** | | **PILFER** | |
|---|---|---|---|---|---|---|---|---|---|
| | | **Computing time [hh:mm:ss]** | **Peak RAM Usage** | **Computing time [hh:mm:ss]** | **Peak RAM Usage** | **Computing time [hh:mm:ss]** | **Peak RAM Usage** | **Computing time [hh:mm:ss]** | **Peak RAM Usage** |
| **Human** | | | | | | | | | |
| | **Testes** | 5:00:40 | 44.672 | 27:21:14 | 18.75 | 1:44:32 | 16.96 | 3:58:59 | 3.58 |
| | **Ovaries** | 0:43:19 | 2.496 | 08:53:54 | 2.88 | 1:24:25 | 16.96 | 2:46:29 | 2.82 |
| | **Somatic** | 0:07:54 | 1.472 | 0:11:27 | 1.472 | 0:57:36 | 7.808 | | 0.81 |
| **Mouse** | | | | | | | | | |
| | **Testes** | 2:37:50 | 9.024 | 26:12:31 | 7.32 | 3:29:04 | 25.02 | 19:03:52 | 4.48 |
| | **Ovaries** | 0:46:41 | 3.776 | 03:53:50 | 1.00 | 1:03:17 | 6.66 | 0:07:46 | 1.344 |
| | **Somatic** | 0:25:58 | 7.712 | 1:34:44 | 4.224 | 0:21:27 | 25.02 | 0:06:54 | 0.32 |
| **Fly** | | | | | | | | | |
| | **Testes** | 2:13:56 | 18.816 | 26:34:16 | 14.212 | 0:23:50 | 9.984 | 23:00:51 | 2.88 |
| | **Ovaries** | 1:45:6 | 7.424 | 14:20:02 | 7.87 | 0:15:40 | 6.784 | 70:23:12 | 2.432 |
| | **Somatic** | 1:07:50 | 7.872 | 23:01:10 | 4.224 | 0:21:27 | 9.856 | 0:26:56 | 4.416 |
